# Supramolecular assembly of the *E. coli* LdcI upon acid stress

**DOI:** 10.1101/2020.05.12.090381

**Authors:** Matthew Jessop, Clarissa Liesche, Jan Felix, Ambroise Desfosses, Megghane Baulard, Virgile Adam, Angélique Fraudeau, Karine Huard, Grégory Effantin, Jean-Philippe Kleman, Maria Bacia-Verloop, Dominique Bourgeois, Irina Gutsche

## Abstract

Pathogenic and commensal bacteria often have to resist the harsh acidity of the host stomach. The inducible lysine decarboxylase LdcI buffers the cytosol and the local extracellular environment to ensure enterobacterial survival at low pH. Here, we investigate the acid-stress response regulation of *E. coli* LdcI by combining biochemical and biophysical characterisation with negative stain and cryo-electron microscopy, and wide-field and super-resolution fluorescence imaging. Due to deleterious effects of fluorescent protein fusions on native LdcI decamers, we opt for three-dimensional localisation of nanobody-labelled endogenous wild-type LdcI in acid-stressed *E. coli* cells, and show that it organises into distinct patches at the cell periphery. Consistent with recent hypotheses that *in vivo* clustering of metabolic enzymes often reflects their polymerisation as a means of stimulus-induced regulation, we show that LdcI assembles into filaments *in vitro* at physiologically relevant low pH. We solve the structures of these filaments and of the LdcI decamer formed at neutral pH by cryo-electron microscopy, and reveal the molecular determinants of LdcI polymerisation, confirmed by mutational analysis. Finally, we propose a model for LdcI function inside the enterobacterial cell, providing a structural and mechanistic basis for further investigation of the role of its supramolecular organisation in the acid stress response.

**Significance statement:** Bacteria possess a sophisticated arsenal of defence mechanisms that allow them to survive in adverse conditions. Adaptation to acid stress and hypoxia is crucial for the enterobacterial transmission in the gastrointestinal tract of their human host. When subjected to low pH, *E. coli* and many other enterobacteria activate a proton-consuming resistance system based on the acid-stress inducible lysine decarboxylase LdcI. Here we develop generally-applicable tools to uncover the spatial localisation of LdcI inside the cell by super-resolution fluorescence microscopy, and investigate the *in vitro* supramolecular organisation of this enzyme by cryo-EM. We build on these results to propose a mechanistic model for LdcI function and offer tools for further *in vivo* investigations.

## Introduction

Cell survival requires the adaptation of metabolism to changing environmental demands. Biochemical regulation of metabolic enzymes by cellular metabolites has been intensely studied for many decades. In addition, a growing body of recent light microscopy observations highlights the spatial regulation of enzymes by stimulus-induced phase separation into distinct loci – liquid droplets, amyloid-like aggregates or ordered polymers (1). In eukaryotes, the observed condensation of fluorescently-labelled metabolic enzymes is triggered by stress, including medium acidification, hypoxia and nutrient limitation. Enterobacteria such as *Escherichia coli*, *Salmonella* and *Vibrio* encounter these types of conditions in the host gastrointestinal tract (2). One of the key enterobacterial proteins expressed during the acid stress response, upon oxygen limitation and regulated by the nutrient stress alarmone guanosine tetraphosphate (ppGpp), is the acid stress-inducible lysine decarboxylase LdcI (3–6). LdcI has been scrutinised since the early 1940s because of direct links between the efficiency of the acid stress response and pathogenicity (7–9). This enzyme transforms lysine into cadaverine while consuming protons and producing CO2, thereby contributing to buffering of the bacterial cytosol and the extracellular medium under acid stress conditions to promote bacterial survival. While both the structure and the function of LdcI have been thoroughly studied (6, 10), nothing is known about its localisation inside the bacterial cell, because its fluorescent protein fusions were previously described to form inclusion bodies (11).

Whereas the overwhelming majority of super-resolution fluorescence imaging is focused on eukaryotes, optical studies of bacterial systems are nearly exclusively centred on large macromolecular complexes with obvious superstructure such as cytoskeletal, cell division, chromosome partitioning, RNA degradation and secretion machineries (12–14). However, the relevance of the documented patchy or long-range helical localisation of some of these assemblies is now questioned and requires re-evaluation. Indeed, the vast majority of these studies were based on labelling with either fluorescent proteins or epitope tags shown to be able to induce artefactual associations and localisations (13–16).

A handful of examples of regulation of bacterial metabolic enzymes by phase separation through stimulus-triggered polymerisation concern well-conserved oligomeric proteins involved in nucleotide and amino acid metabolism, such as CTP synthase (17, 18) and glutamine synthetase (19). Interestingly, these enzymes are also able to assemble into filaments *in vitro*, and their *in vivo* condensates, detected both in bacteria and in eukaryotes, have been suggested to correspond to the polymerised state of the enzymes. Other examples of bacterial metabolic enzymes purified as polymers from bacterial extracts or forming polymers *in vitro* are the aldehyde-alcohol dehydrogenase AdhE (20) and the hydrogen-dependent CO_2_ reductase HDCR (21).

Specific to bacteria, LdcI is a decamer composed of five dimers tightly arranged into pentameric double-rings (PDB ID: 3N75) (6). Interestingly, *in vitro* these double-rings were observed by negative stain electron microscopy (ns-EM) to stack on top of one another into filament-like structures (6, 8) but only at non-physiological pH below 5, under which LdcI should not be significantly expressed. Here, to address the spatial localisation of the *E. coli* LdcI, we start by critically evaluating labelling artefacts with an ambition to define the optimal constructs for subsequent chromosomal manipulations. To this end, we overexpress and purify different fluorescent protein (FP) fusions of LdcI, perform their structural and biochemical characterisation *in vitro*, and observe the distribution of the overexpressed constructs *in vivo*. This methodological section enables us to propose a workflow that brings together examinations of *in vivo* protein localisations with *in vitro* biochemical and ns-EM characterisations of the purified FP fusions in order to ensure artefact-free optical imaging investigations. This analysis is followed by unveiling a patchy distribution of endogenous LdcI inside the *E. coli* cell upon acid stress. Because this condensation of LdcI into patches is reminiscent of stress-induced polymerisation, in particular in yeast, we investigate the *in vitro* architecture of LdcI at pH 5.7. We show that at this physiological pH, corresponding to the optimum of the LdcI enzymatic activity, LdcI is a polymer assembled of stacked decamers, and solve the polymer structure by cryo-electron microscopy (cryo-EM). For comparison, we also determined the cryo-EM structure of the LdcI decamer at neutral pH. In addition, mutational analysis of the LdcI stack-forming interfaces allowed identification of critical residues involved in stack formation. Finally, we discuss the observed LdcI localisation pattern in the light of the wealth of available functional and imaging data, and offer a structural and mechanistic basis of supramolecular LdcI assembly, which will aid in the design of future experiments linking LdcI stack formation to *E. coli* acid stress fitness.

## Results

### Fluorescent protein fusions affect LdcI structure without modifying localisation of the overexpressed fusion constructs

Because of the small size of the *E. coli* cell, we opted for super-resolution microscopy imaging (22) and set out to localise LdcI inside the cell upon acid stress by either Photoactivation Localisation Microscopy (PALM) or Stochastic Optical Reconstruction Microscopy (STORM) (23–25). *A priori*, PALM seemed more relevant because this technique relies on genetically-encoded FPs fused to the protein of interest, and is therefore unmatched in terms of labelling specificity and efficiency. In addition, PALM does not require delivery of fluorescent molecules across the cell wall, and enables live cell imaging and single molecule tracking.

Considering that LdcI is an acid stress response enzyme, we opted for either mGeosM (26) or Dendra2_T69A_ (27) as FP markers because of their relatively low pKa values, high monomericity and high fluorescent quantum yields. In the LdcI decamer structure, the N-termini are oriented inwards and towards the central pore of the double ring, while the C-termini point to the ring periphery and are readily accessible from the outside. Therefore, on the one side, one could assume that an N-terminal labelling with an FP would be likely to interrupt the LdcI tertiary structure. On the other side, the only well characterised binding partner of LdcI, the AAA+ ATPase RavA, is known to interact precisely with the C-terminal β-sheet of LdcI (28). Assembly of two copies of double-pentameric rings of LdcI and five copies of hexameric RavA spirals results in a huge 3.3 MDa macromolecular cage of intricate architecture and largely unknown function (28–30). The LdcI-RavA cage is proposed to assist assembly of specific respiratory complexes in *E. coli* and to counteract acid stress under starvation conditions by alleviating ppGpp-dependent inhibition of LdcI (31–33). Thus, although these functions still require further investigation, preservation of the RavA-binding propensity should be one of the criteria for assessing the suitability of an LdcI-FP fusion. Therefore, both N- and C-terminal fusion constructs with either mGeosM or Dendra2_T69A_ attached to LdcI via an appropriate linker were cloned into dedicated plasmids and overexpressed at conditions optimised for LdcI overproduction (see Methods for details). Expression of fusion proteins was immediately detected by wide-field fluorescence imaging that showed a similar distribution for the four fusions (Figure 1A). Each construct was then purified in order to assess its structural integrity and RavA binding capacity *in vitro*, with a goal of defining the most suitable construct for the subsequent creation of a corresponding chromosomal fusion.

**Figure 1.**
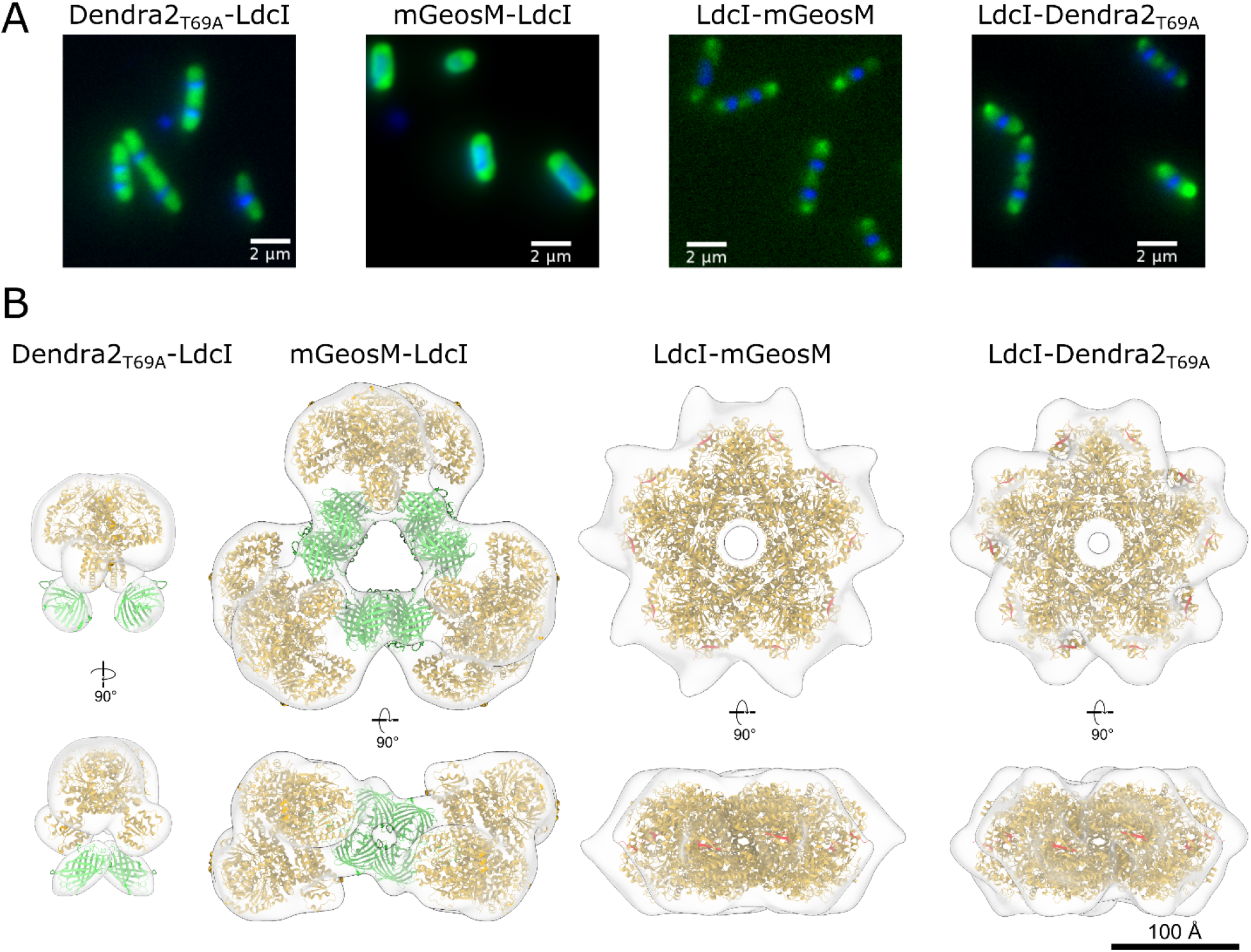
Fluorescent protein fusions affect LdcI structure without significantly altering cellular localisation of overexpressed constructs. **A)** Wide-field fluorescence microscopy images of *E. coli* cells overexpressing the fluorescent fusion proteins in A). Green fluorescence = FP-LdcI, blue fluorescence = DAPI-stained DNA. **B)** ns-EM 3D maps of fluorescently-tagged LdcI, with fitted models. Left – Dendra2_T69A_-LdcI forms dimers, with fluorescent barrels located next to the N-terminus of LdcI as expected. Second from left – mGeos-M-LdcI forms large non-native oligomers, composed of three LdcI tetramers bridged by tetramers of mGeosM. Second from right and right – both C-terminal fluorescent fusions LdcI-mGeosM and LdcI-Dendra2_T69A_ form decamers, with protrusions at the C-terminus (coloured in red) attributed to flexibly-linked fluorescent proteins. Fitted PBD models are as follows – LdcI: 3N75; Dendra2: 2VZX; mGeosM: 3S05 (mEos2 crystal structure).

The N-terminal Dendra2_T69A_-LdcI fusion formed exclusively dimers, confirming the structure-based prediction that the fluorescent tag at this position would perturb the dimer-dimer interaction (Figure 1B, Supplementary Figure 1). Admittedly, native LdcI was shown to dissociate into dimers *in vitro* at pH above approximately 7.5, but in this pH range LdcI is not supposed to be expressed in the cell and therefore this dissociation may be irrelevant. Surprisingly, in contrast to Dendra2_T69A_-LdcI, mGeosM-LdcI assembled into regular symmetric non-native higher-order oligomers, with a dramatically altered quaternary structure (Figure 1B, Supplementary Figure 1). These oligomers were built of three LdcI tetramers, bridged together by additional densities. Noteworthy, mEos2, the fluorescent protein from which mGeosM was derived, crystallises as a tetramer (PDB ID: 3S05) that can be straightforwardly fitted into the LdcI-bridging densities (Figure 1B). This illustrates that despite the fact that mGeosM was designed as a monomeric FP (with the first “m” explicitly standing for monomeric), some residual oligomeric tendency is still maintained. This propensity of mGeosM to oligomerise when bound to LdcI may be driven by avidity effects – as LdcI dimers begin to assemble into a decamer, the local concentration of mGeosM increases to such a point that oligomerisation becomes energetically favourable, despite the apparent monomer behaviour of mGeosM in gel filtration (26). This is also in line with the known propensity of mEos2, from which mGeosM is derived, to form tetramers at high concentration (34). To conclude, both N-terminal fusions induced non-native assembly of LdcI, and therefore neither were appropriate for determining the native localisation of this enzyme in the cell.

As expected, C-terminal LdcI fluorescent fusions formed decamers with protruding densities that can be attributed to flexibly attached FPs (Figure 1B, Supplementary Figure 1). Nevertheless, despite the long flexible linker between LdcI and the FP, these constructs were unable to interact with RavA as shown by Bio-Layer Interferometry (BLI) binding studies (see Methods and Supplementary Figure 1 for details). This means that the functionality of these fusions cannot be considered as entirely retained. Thus, none of the four fusions were suitable for an in-depth imaging analysis under conditions of native LdcI expression upon acid stress that we planned to undertake. Altogether, we demonstrated that structural integrity and unaltered interaction with known partners are useful read-outs for functional preservation. Based on this result we propose that, when feasible, purification and structure-function analysis of FP fusions should be performed prior to interpretation of the protein localisation inferred from observation of FP fusions by optical methods. This workflow can be used for example in cases where chromosomal manipulation for assessment of intact function is difficult and/or the phenotype is condition-dependent or unclear, whereas basic *in vitro* biochemical and ns-EM characterisation are efficiently set up.

### Immunofluorescence of LdcI in *E. coli* under acid stress reveals its supramolecular organisation

Because none of the FP fusions possessed the properties of native LdcI, we decided to turn to STORM imaging of the endogenous enzyme. One of the drawbacks of STORM is the requirement for exogenous labelling and therefore the fixation and permeabilisation of cells. These procedures are notoriously known to potentially affect cell morphology, and therefore, when possible, the usage of live cell imaging with FP fusions (as for example in PALM), or observation of unfixed samples under cryogenic conditions would be ideal. Moreover, fixation, permeabilisation and specific exogenous labelling in bacteria present unique challenges because of the complex, often multi-layered cell wall (35, 36), A significant advantage of STORM however is the possibility of direct imaging of the wild-type (WT) protein, which should circumvent the dangers associated with FP fusions described above. The three prerequisites for imaging of WT systems by immuno-based labelling in general and of LdcI in particular are (i) availability of an antibody or nanobody (antibody fragment derived from heavy-chain-only camelid antibody) coupled to an organic fluorescent dye and directed towards the native protein, (ii) precise knowledge of endogenous expression conditions, (iii) validation of an efficient permeabilisation and immunolabelling technique that enables the antibody/nanobody to enter the cell and specifically target the protein of interest without altering the native organization of the cell ultrastructures.

As a first step towards STORM imaging of endogenous LdcI, we probed an anti-LdcI nanobody, hereafter called anti-LdcI-Nb. The complex between purified *E. coli* LdcI and the nanobody was purified by size exclusion chromatography (see Methods), and imaged by ns-EM. The resulting 3D map of the LdcI/anti-LdcI-Nb complex demonstrates binding of anti-LdcI-Nb to each LdcI monomer in the decamer and reveals the location of the interaction site which is clearly distinct from the RavA binding site (Figure 2A, Supplementary Figure 2). This nanobody was thus identified as a suitable labelling agent for LdcI imaging in *E. coli* cells, and labelled with the fluorescent dye AlexaFluor647 (AF647) or AlexaFluor488 (AF488) (see Methods).

**Figure 2.**
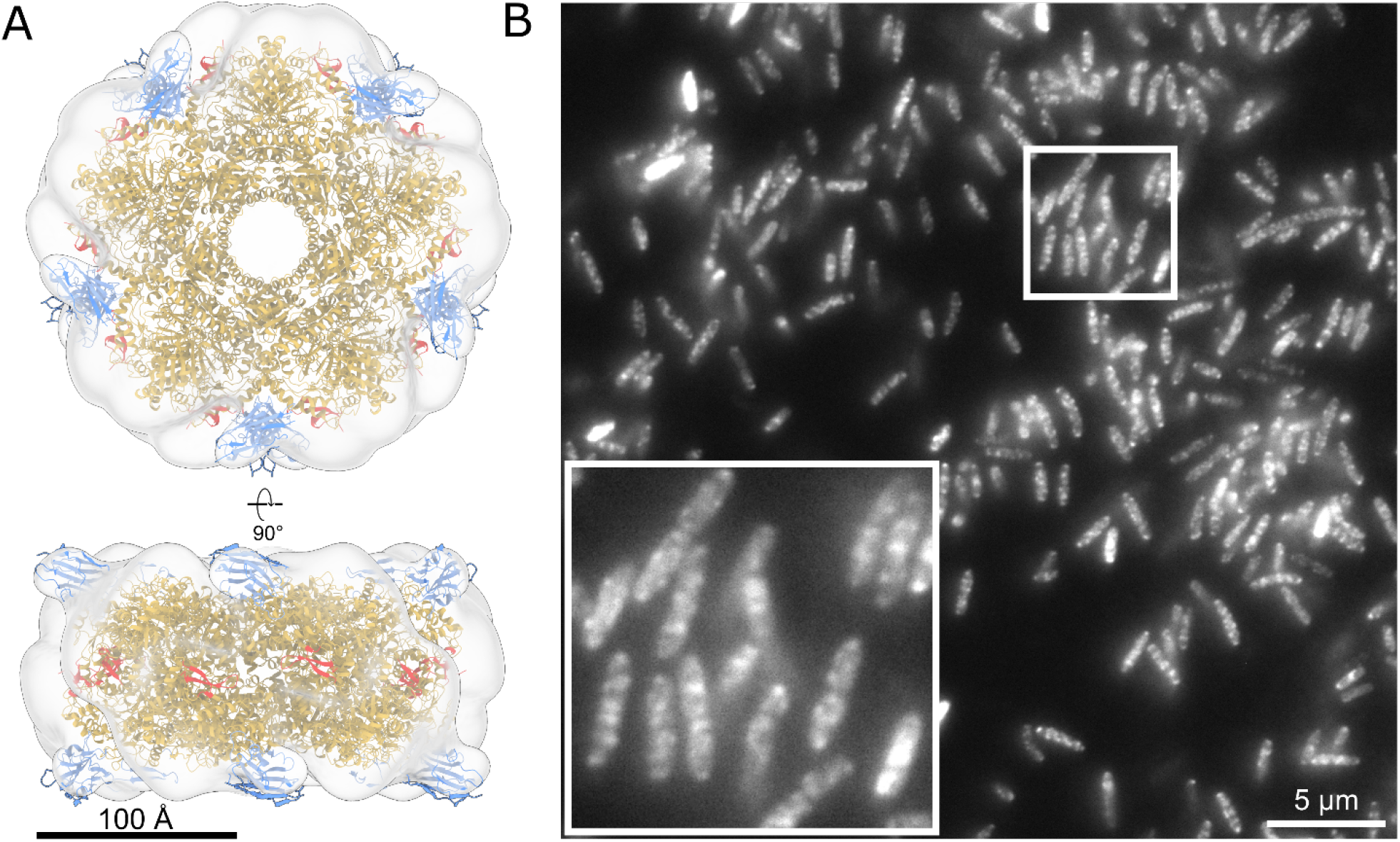
Anti-LdcI-Nb is a useful tool to probe cellular localisation of endogenous LdcI under acid stress conditions. **A)** Ns-EM 3D reconstruction of the LdcI decorated by anti-LdcI-Nb. An LdcI decamer (gold, PDB ID: 3N75) and 10 nanobodies (blue, PDB ID: 1MEL, anti-lysozyme nanobody) are fitted in the density, with the nanobody binding at the top and bottom of the decameric ring. The C-terminal RavA binding site is indicated in red, and is in a spatially distinct location from the bound nanobodies. Scale bar = 100Å. **B)** Wide-field fluorescence microscopy image of wild-type *E. coli* MG1655 cells grown at pH 4.6 for 90 minutes stained with anti-LdcI-Nb labelled with AF488. Inset – zoom of the image showing punctuate fluorescence patterns. Scale bar = 5 μm.

Consistently with published data, LdcI expression could be induced by a pH shift experiment (i.e. transfer of bacteria from pH 7.0 into a pH 4.6 growth medium) in the presence of lysine under oxygen-limiting conditions (Supplementary Figure 3A). While the WT strain grew well under these conditions and efficiently buffered the extracellular medium up to pH 6.2 in approximately 1.5 to two hours (Supplementary Figure 3A) concomitantly with the increase in the level of LdcI expression (Supplementary Figure 3B, C), the growth and the acid stress response of the Δ*ldcI* mutant strain were severely impaired (Supplementary Figure 3A). Since the peak of LdcI expression by the wild type cells was achieved between one and two hours after exposure to acid stress (Supplementary Figure 3B, C), a time point of 90 minutes was chosen for the subsequent labelling and imaging experiments. The specificity and performance of anti-LdcI-Nb in immunofluorescence labelling of permeabilised *E. coli* cells were characterised by flow cytometry and wide-field fluorescence imaging (Supplementary Figure 3D, E). Both techniques demonstrated that in the absence of LdcI expression no specific fluorescence is seen, confirming the suitability of anti-LdcI-Nb for immunofluorescence studies. Thus, the above-mentioned prerequisites for immuno-based imaging of cellular LdcI have been fulfilled. Noteworthy, the LdcI expression profile highlighted a considerable asset of the usage of STORM instead of PALM for visualisation of the endogenously expressed LdcI: indeed, the transient nature of the expression and the necessity of work under oxygen-limiting conditions may have created difficulties due to the longer maturation time of the FPs under these conditions.

Initial characterisation of the cellular distribution of LdcI 90 minutes after exposure of *E. coli* cells to acid stress was carried out by wide-field fluorescence imaging. Based on these images, it appeared that natively expressed LdcI did not display a homogeneous cytoplasmic distribution but rather showed a patchy localisation pattern (Figure 2B). 3D STORM imaging subsequently provided a more detailed view of this patchy distribution. As shown in Figure 3, Supplementary Figure 4, and Supplementary Movies 1-5, the labelling density was lower in the centre of the bacterium. This indicates a propensity for LdcI to cluster near the cell periphery, towards the inner membrane and the cell poles, rather than being distributed homogeneously through the volume. Recent investigations argue that, in most cases, *in vivo* clustering of metabolic enzymes corresponds to their polymerised states, and represents an efficient means of regulation of enzymatic activity and metabolic homeostasis in response to a stimulus (1). Thus, structure determination of these polymers is a crucial step towards the understanding of regulation mechanisms. We previously documented that *in vitro* at pH below 5 and high concentration, LdcI decamers indeed tend to stack (6). However, at pH 5.7, optimal for LdcI expression and enzymatic activity and likely corresponding to the internal pH upon acid stress, the oligomeric state of LdcI has not yet been addressed by EM. Supposing that the high local concentration of LdcI, clustered in patches in the *E. coli* cell under acid stress conditions, is likely to induce enzyme polymerisation via stack formation, we next examined the LdcI assembly state at pH 5.7 by cryo-EM and discovered that it does indeed form straight polymers (Figure 4).

**Figure 3.**
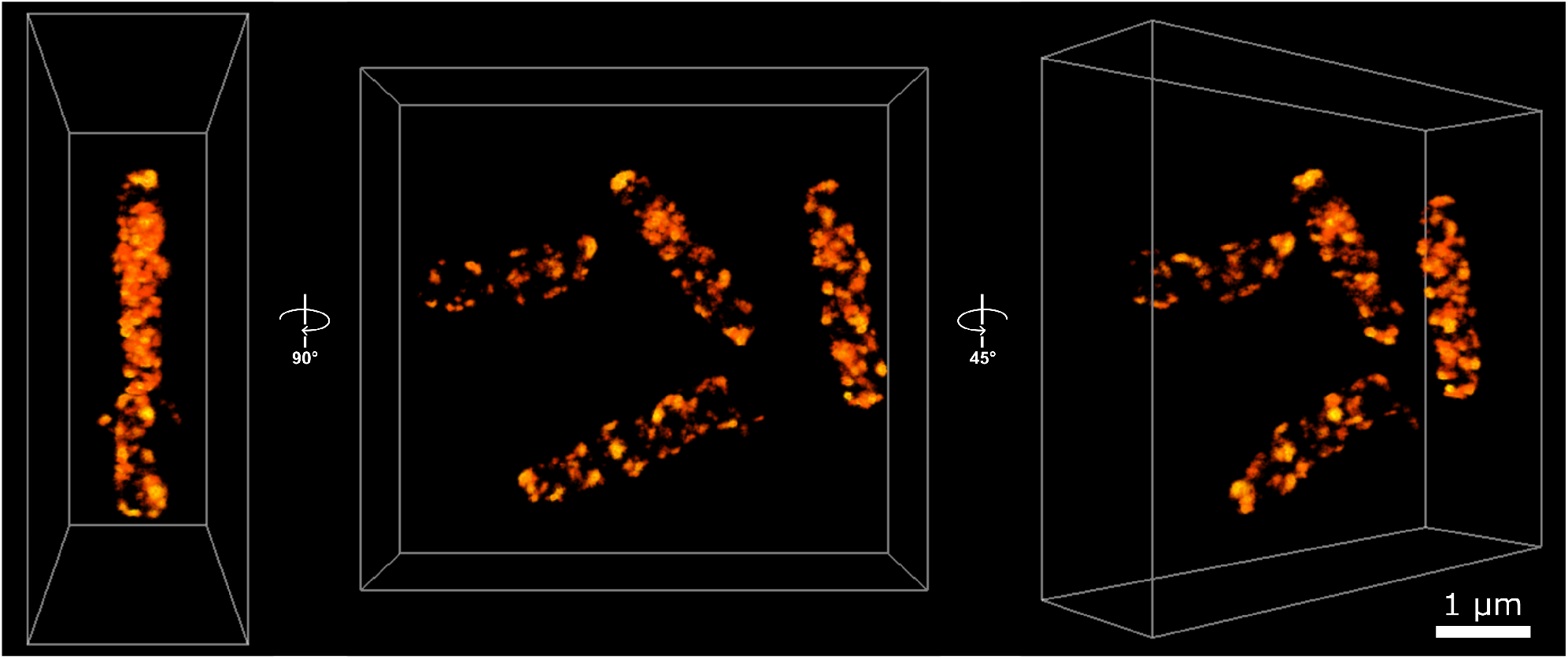
3D STORM imaging of *E. coli* cells stained with anti-LdcI-Nb reveal a patchy pseudo-helically arranged distribution of endogenous LdcI upon acid stress. 3D-STORM imaging of native LdcI in wild-type *E. coli* cells 90 minutes after exposure to acid stress (see Methods) with AF647-conjugated anti-LdcI-Nb. Points are coloured according to localisation density, with brighter points corresponding to higher localisation density. The centre panel shows four cells in the field of view, looking down the z-axis. Left and right panels show side and tilted views respectively. Scale bar = 1 nm.

**Figure 4.**
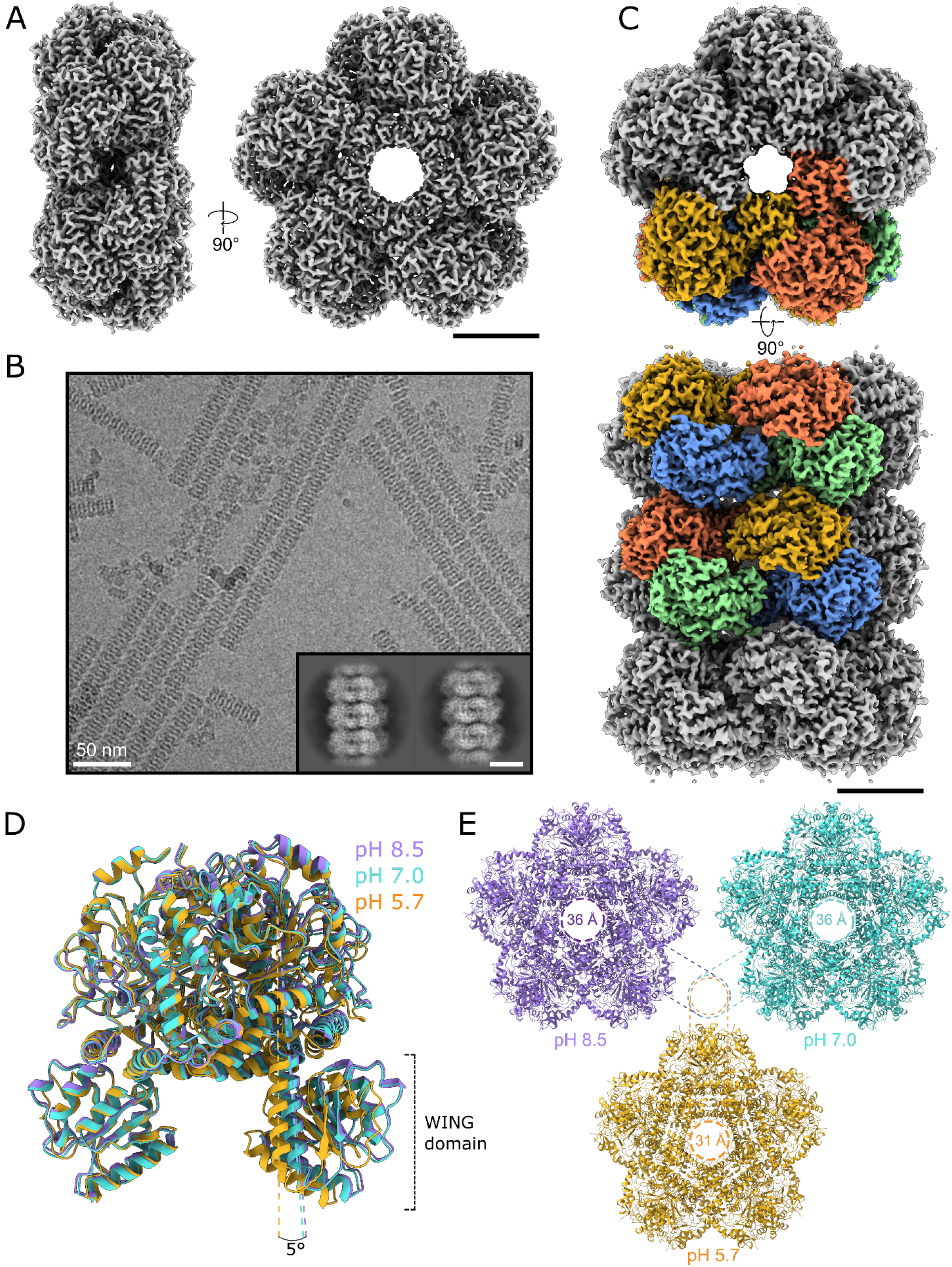
Cryo-EM analysis of the LdcI decamer at neutral pH and LdcI stacks formed in acid stress conditions. **A)** Cryo-EM reconstruction of the LdcI decamer at pH 7.0 from side (left) and top (right) views. Scale bar = 50 Å. **B)** Cryo-EM micrograph of LdcI stacks at pH 5.7, scale bar = 100 nm. Inset – 2D class averages displaying clear secondary structural features, scale bar = 100 Å. **C)** Top (above) and side (below) views of the cryo-EM reconstruction of a three-decamer LdcI stack. Four dimers are coloured either blue/gold or green/coral, corresponding to the colouring of the atomic model presented in Figure 5. Scale bar = 50 Å. **D)** Overlay of LdcI dimers at pH 8.5 (PDB ID: 3N75, shown in lilac), 7.0 (shown in cyan) and 5.7 (shown in gold). Alignment was carried out on a single monomer in the dimer pair. There is a 5° shift in the angle between the wing domains of LdcI at pH 5.7 and pH 7.0/8.5. **E)** Comparison of the central decamer pore diameter between LdcI at pH 8.5, 7.0 and 5.7, showing a 5 Å decrease in the pore size upon stack formation at low pH.

### Structural determinants of LdcI stack formation revealed by cryo-EM

In order to understand the molecular determinants of LdcI polymerisation at low pH and to provide a framework for the analysis of LdcI function under acid stress, we solved the 3D structures of the LdcI decamer at pH 7.0 and of LdcI stacks at pH 5.7 by cryo-EM (See Methods, Figure 4, Supplementary Figures 5-6, Supplementary Table 1). The 2.8 Å resolution structure of the LdcI decamer at pH 7.0 (Figure 4A) is extremely similar (Supplementary Table 2 & 3) to the LdcI crystal structure solved at pH 8.5 in an inhibited ppGpp-bound state (6). However, contrary to pH 7.0 and even pH 6.2 where LdcI is still predominantly decameric (10), LdcI forms straight rigid filaments on the cryo-EM grid at pH 5.7, which corresponds to the pH of maximum LdcI enzymatic activity (Figure 4B). The structure of a three-decamer stack was solved to a resolution of 3.3 Å, revealing the structural details of acid stress-induced LdcI polymerisation (Figure 4C). LdcI decamers stack tightly on top of one other, with negligible rotation between decamers along the stack. Each dimer fits snugly in the inter-dimer groove of decamers above and below.

A comparison of LdcI decamer structures taken from the LdcI stack cryo-EM map (at pH 5.7), the LdcI decamer cryo-EM map (at pH 7.0) and the crystal structure of decameric LdcI crystallised with bound ppGpp at pH 8.5 (PDB ID: 3N75) reveals some remarkable differences between the stack structure and the two decamer structures (Figure 4D, E, Supplementary Table 2 & 3). While the three structures do not show any major differences at the monomer level, a structural alignment of an LdcI dimer extracted from the LdcI decamer structures with a dimer extracted from the LdcI stack structure uncovers a rigid body-like rotation between monomers around a hinge region located at the monomer-to-monomer interface (Supplementary Table 2 & 3, Figure 4DE). This rotation results in a 5° tilt when comparing the N-terminal wing-domains in LdcI dimers, and an overall slightly decreased diameter of the central cavity inside the stacked LdcI rings (Figure 4D, E), which may contribute to the tight packing of each dimer into the grooves of an opposing decamer in the stack.

A careful examination of the LdcI stack structure shows that two major inter-decamer interfaces situated at a two-fold symmetry axis perpendicular to the stack direction contribute to stack formation (Figure 5A). In particular, the first interface (Figure 5B) is formed between residues K422, D460, R468, D470, and E482 situated in the ppGpp-binding domain (amino acids 418-564), and residues N314, D316, and G352 from the PLP-binding domain (amino acids 184-417). D460 from one decamer makes an electrostatic interaction with K422’ of a neighbouring decamer in the stack. R468 is sandwiched between D316’ and E344’ from a neighbouring decamer, and makes electrostatic interactions through its η1 and η2 nitrogen atoms with D316’. In addition, D470 interacts with the backbone of G352’, and E482 forms hydrogen bonds with N314’. The second interface (Figure 5B) is formed between residue N94 of the wing domain (amino acids 1-130) of one set of opposing dimers, and a stretch of four residues in the ppGpp-binding domain – T444, E445, S446 and D447 - at the end of helix α16 from a second set of opposing dimers. The wing domain residue N94 makes hydrogen bonds with E445’ of an opposing dimer. A second charged residue, D447, interacts with the backbone of T444’, and is held in place by R97 from the wing domain of a neighbouring dimer in the same decamer.

**Figure 5.**
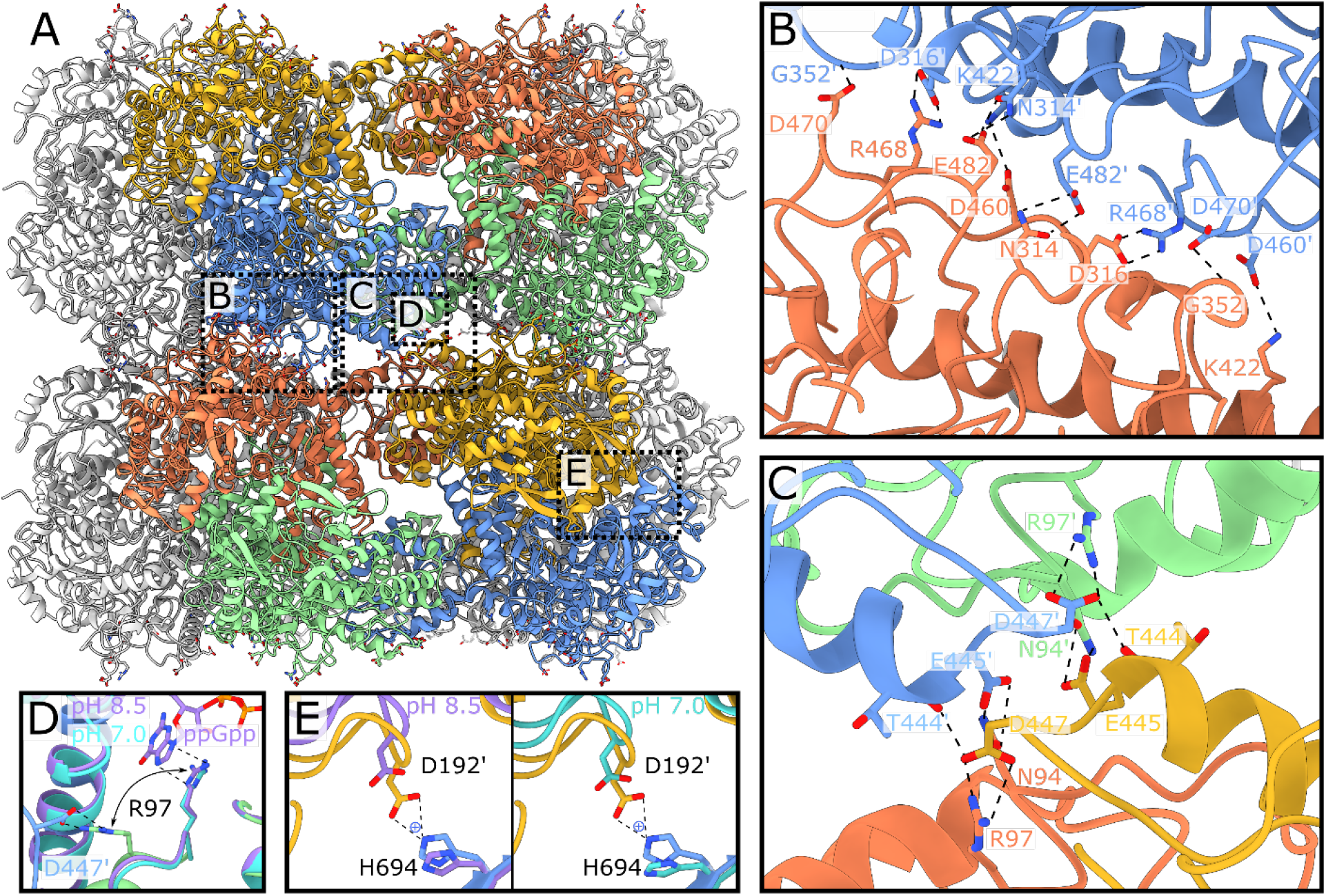
Structural insights into molecular determinants of the LdcI polymerisation under acid stress conditions. **A)** Atomic model of a two-decamer LdcI stack at pH 5.7. Dimers are coloured as shown in the cryo-EM map in Figure 4. Dotted boxes on the two-decamer stack indicate the locations of the zooms shown in panels B-E. **B)** Close-up of the first decamer-decamer interface, which includes the key stack-forming residue R468. **C)** Close-up of the second decamer-decamer interface. **D)** Overlay of the LdcI decamer structures at pH 8.5 and 7.0 with the LdcI stack structure at pH 5.7, focussed on R97. R97 in the LdcI stack (green) adopts a different conformation compared to the one in the pH 8.5 crystal structure (purple, with ppGpp bound) and the pH 7.0 cryo-EM map (cyan, without ppGpp bound). Despite the absence of ppGpp in the pH 7.0 sample, R97 is still oriented towards the ppGpp binding site. **E)** Comparison between the H694-D192’ distance in the LdcI stack at pH 5.7 (coloured gold and blue), the LdcI decamer at pH 8.5 (left, coloured purple) and pH 7.0 (right, coloured cyan). Key residues are labelled for all panels, and interactions are shown with dotted lines.

Considering that the LdcI polymerisation is induced by acid stress, we wondered which residues in the interface would be sensitive to pH changes. Surprisingly, most of the side chains involved in the inter-decamer interface are charged arginine (pKa 13), aspartate and glutamate residues (pKa of 4 and 3 respectively), which do not change protonation state in the pH window relevant for LdcI activity (pH 5-7) (37). Nonetheless, other residues, situated outside the interaction interfaces, may drive stack formation through pH-dependent interactions that would in turn lead to the observed inter-monomer rotation and the associated constriction of the LdcI central cavity, coupled to the alignment of complementary contacts at interfaces one and two (Figure 4D, E). We note for example that H694 should be protonated in the LdcI stack structure at pH 5.7 but deprotonated in the two ring structures at pH 7.0 and pH 8.5, and that an electrostatic interaction between H694 and D192 situated in the linker region is present in the stack structure only (Figure 5E).

To validate the observed interactions at the inter-decamer interface, and to assess the individual importance of key residues involved in LdcI stack formation, we constructed four LdcI point mutants (R468E, R97E, H694A and H694N), two double mutants (R97E/R468E and E445A/D447A) and one triple mutant (E445A/D447A/R468E). The mutants were purified following the protocol for wild-type LdcI (see Methods), diluted into a buffer at pH 5.7 and observed by ns-EM (Figure 6, Supplementary Figure 7). Although the grid preparation procedure for ns-EM yields stacks that are shorter and more curved and distorted when compared to the cryo-EM data (Figure 4B, Figure 6), our previous observations of the five paralogous *E. coli* amino acid decarboxylases justify the validity of this approach for a qualitative comparative analysis (6, 37). Ns-EM images make immediately apparent that the E445A/D447A double mutant does not have a significantly altered capability of stack formation at pH 5.7 when compared to WT LdcI, whereas the single R468E mutation is sufficient to completely abolish stack formation. The R97E mutant has a moderate destabilising effect and displays fewer and smaller stacks then the WT LdcI. Consistently with the major role of R468 in the LdcI stack formation, the R97E/R468E double mutant and E445A/D447A/R468E triple mutant exclusively occur as decamers at low pH. Altogether, these results reveal that R468 is one of the key determinants of LdcI stack formation. Finally, similarly to the R97E mutant, a modest destabilisation of stack formation is observed for the two histidine mutants, H694A and H694N, favouring our hypothesis that H694 may have an influence on the propensity of LdcI polymerisation at low pH (Figure 6, Supplementary Figure 7).

**Figure 6.**
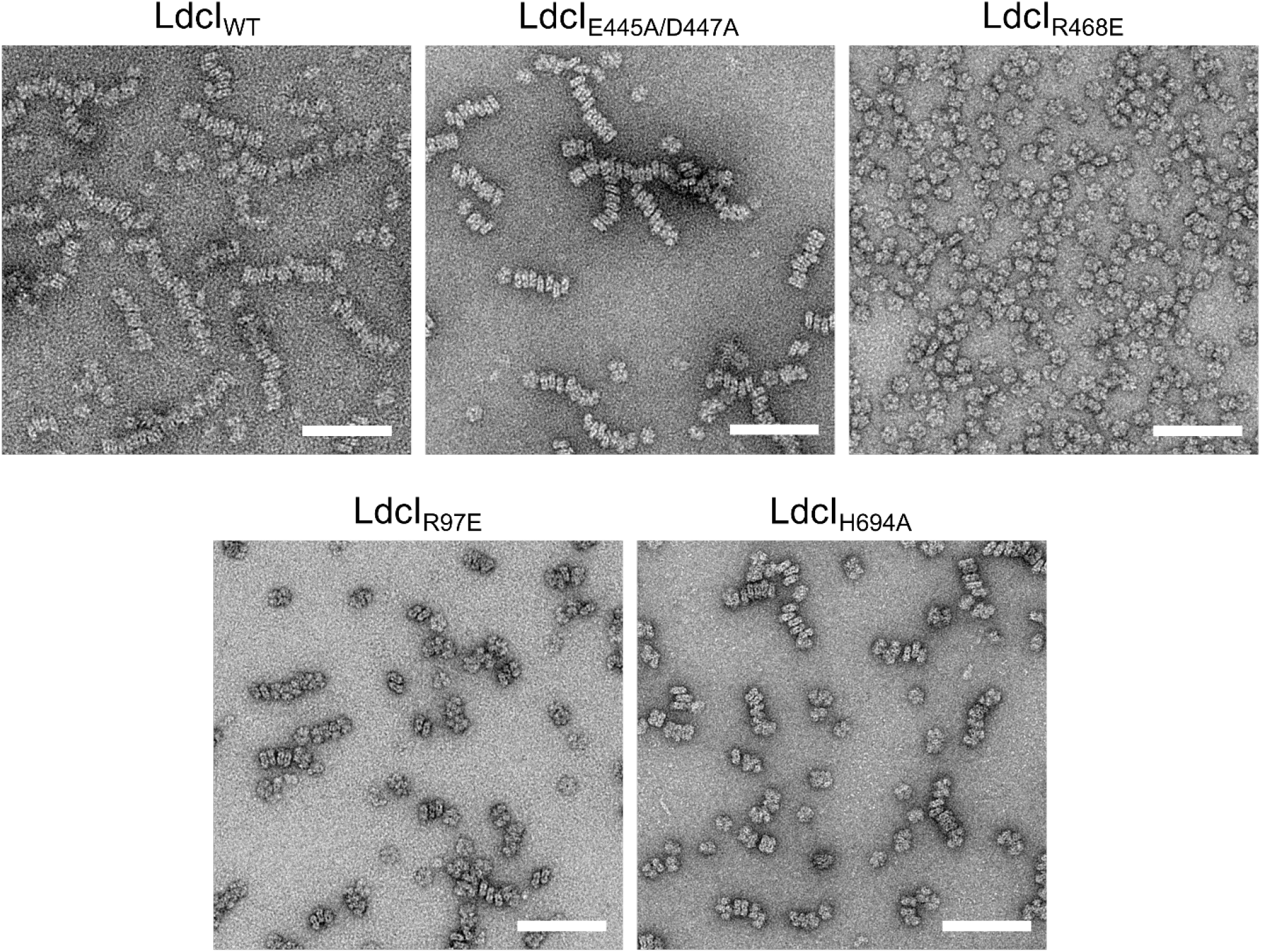
Mutational analysis of the predicted molecular determinants of the LdcI polymerisation under acid stress conditions. Cropped negative stain EM micrographs of wild-type and mutant LdcI at pH 5.7, scale bar = 100 nm. LdcI_WT_ polymerises at pH 5.7, as does the double mutant LdcI_E445A/D447A_. In contrast, the single mutation R468E abolishes stack formation completely. Both LdcI_R97E_ and LdcI_H964A_ are able to polymerise, however the stacks tend to be shorter than for LdcI_WT_. Micrographs for LdcI_R97E/R468E_, LdcI_E445A/D447A/R468E_, and LdcI_H694N_ were also collected but are not shown here. LdcI_E445A/D447A/R468E_ and LdcI_R97E/R468E_ behaved like LdcI_R468E_ and remained entirely decameric at low pH, while LdcI_H694N_ displayed similar behaviour to LdcI_H694A_.

## Discussion

Our synergistic approach, combining several *in vitro* techniques including biochemical characterisation of purified fluorescent protein fusions, ns-EM observation of mutants, low resolution ns-EM reconstruction and high resolution cryo-EM analysis, with *in vivo* flow cytometry, wide-field and 3D-STORM imaging, provides insights into the supramolecular LdcI assembly upon acid stress. This work adds to the very few known examples of regular polymerisation as means of regulation of enzymes involved in amino acid metabolism in bacteria. The cryo-EM structure of the LdcI stacks presented here offers a molecular framework for future investigation of the role of the LdcI polymerisation in the acid stress response.

The high resolution cryo-EM structures of the stack at pH 5.7 and of the decamer at pH 7.0 complement the previously solved crystal structure of the ppGpp-bound LdcI decamer obtained at pH 8.5. Previous serendipitous co-crystallisation of LdcI with the nutrient stress response alarmone ppGpp led to assessment of the effect of ppGpp on LdcI activity, and to a proposal that ppGpp would act as an LdcI inhibitor that prevents excessive lysine consumption upon nutrient limitation during acid stress (6). In addition, a ppGpp-dependent disassembly of the LdcI stacks had been previously observed but could not be structurally explained since the ppGpp binding site is situated between two neighbouring dimers inside the LdcI ring (6). Our cryo-EM structures show that one of the residues involved both in ppGpp binding and in the stack formation is arginine 97 (R97). In the crystal structure of ppGpp-bound LdcI, R97 makes a stacking interaction with the guanosine imidazole ring of ppGpp, while in LdcI stacks R97 is involved in a key interaction at the second interface (Figure 5D), where it locks D447 in a conformation allowing interactions between helices α16 from opposing LdcI decamers. Binding of ppGpp to LdcI interferes with the R97-D447 interaction, thereby most likely prohibiting correct positioning of D447 at the tip of helix α16, and resulting in a disruption of the second stack interface, leading to a moderate yet notable stack destabilisation (Figure 6, Supplementary Figure 7). Furthermore, our cryo-EM structure of ppGpp-free LdcI decamers at neutral pH enables to rule out the effect of ppGpp on the differences observed between the LdcI stack structure at pH 5.7 and the ppGpp-LdcI crystal structure at pH 8.5. Indeed, despite the absence of ppGpp, R97 is still oriented towards the ppGpp binding site and away from the inter-decamer interface in the pH 7.0 decamer cryo-EM map (Figure 5D). This suggests that the conformational changes in LdcI driving stack formation are mostly driven by low pH and not by the absence of ppGpp, although the D192 and H694 hinge residues are similarly far apart in the pH 8.5 and pH 7.0 structures (Figure 5E). Our current work provides a solid experimental and structural basis for a future closer evaluation of the hypothesised role of ppGpp in LdcI regulation *in vitro* and *in vivo (6)*

Although the optical imaging experiments performed in this study do not allow quantitative evaluation of the nature of the clusters of endogenous LdcI at the bacterial periphery, they would presumably correspond to LdcI assembled into stacks. Possible reasons for such assembly may be to provide an efficient way to locally increase the LdcI concentration and to enhance its activity. But why would LdcI, an apparently highly soluble protein, be driven towards the inner membrane? What would be the advantage for acid-stressed *E. coli* cells to increase the concentration of LdcI via stack formation in these particular peripheral locations? Localisation of proteins to specific sites in the bacterial membrane was shown to be generally driven by chemical factors such as the phospholipid composition of the lipid microdomains, and by physical factors such as the degree of local curvature or the electric potential of the membrane (38). An attractive hypothesis would be that as an acid stress response protein performing a proton-consuming enzymatic reaction, LdcI may be attracted to proton sinks formed by anionic phospholipids which compartmentalise oxidative phosphorylation (OXPHOS) complexes for efficient functioning in bacterial respiration and adaptation to environmental changes. Indeed, OXPHOS complexes were often described to be unevenly distributed in the membrane in the form of mobile patches (39–43), providing evidence for highly dynamic and spatially organised bioenergetic membranes in *E. coli* cells (43). In addition, certain bacterial flottilins, which are essential scaffold proteins of the functional membrane microdomains, equivalent to the lipid rafts of eukaryotic cells, also show a patchy distribution and were shown to interact with specific OXPHOS complexes (43, 44). In this regard, two different lines of evidence would be interesting to note. First, LdcI was described to co-purify with a partially assembled Complex I (45), whereas the LdcI-binding partner RavA, as well as ViaA, the second protein expressed from the *ravAviaA* operon and which also interacts with RavA, were shown to interact with both Complex I and fumarate reductase (32, 33). Second, the other *E. coli* PLP-dependent lysine decarboxylase LdcC, exercising the role of cadaverine biosynthesis irrespectively of acid stress (46), neither binds RavA nor forms stacks (29, 37), in spite of its 69% of identity with LdcI. It is interesting to note that in LdcC the key stack-forming residue R468 has been substituted for an alanine while the interacting D316 has been preserved.

Our STORM images suggest that individual patches have a peripheral distribution with a long-range stripy or pseudo-helical organisation (Figure 3, Supplementary Figure 4, Supplementary Movies 1-5). While similar distributions have been documented for bacterial cytoskeletal, cell division, chromosome partitioning, RNA degradation and secretion machineries, an eventual impact of labelling on these distributions, demonstrated specifically for the YFP-MreB (47), warrants caution in their interpretation (13, 15). Here however, we observed endogenous wild-type LdcI in cells fixed prior to their permeabilisation and labelled with anti-LdcI-Nb, which means that the resulting pattern is likely real. In addition, examination of some published images of OXPHOS patches (for example Llorente-Garcia et al., 2014) also hints to a similar organisation. Excitingly, anionic phospholipid-specific dyes and fluorescently-labelled antibiotics specific for nascent peptidoglycan synthesis upon cell elongation were also proposed to be distributed on helical or stripe patterns (38, 48–51). It is therefore tempting to imagine that inside the bacterial cell, LdcI has a tendency to follow a general path upon polymerisation governed by the underlying chiral patterns in the cell envelope (52).

Finally, from the methodological view, our work convincingly illustrates that different FP fusion constructs can share the same cellular distribution in spite of a completely different structure, necessitating caution when inferring intact function from the preservation of the protein localisation inside the cell. Our findings emphasize the importance of characterising FP fusions using both biochemical and structural techniques, such as ns-EM, to ensure that the FP tag disrupts neither structure nor function of the target protein.

## Methods

### Expression constructs

For fluorescence studies, several FP fusion constructs were generated starting from an available plasmid containing the coding sequence of LdcI (Uniprot entry P0A9H3), cloned in the pET22b(+) vector with a C-terminal 6xHis-tag (6). All constructs were generated using the Gibson cloning strategy and verified by sequencing analysis. The Gibson assembly was performed using 0,4U T5 exonuclease, 2,5U Phusion polymerase and 400U Taq ligase (New England Biolabs) in 1X ISO buffer (100mM Tris-HCl pH 7,5, 10 mM MgCl2, 0.8 mM dNTP mix, 10mM DTT, 50 mg PEG-8000, 1 mM NAD). 7.5 μL of the GIBSON Master Mix was mixed with 2.5 μL DNA, containing circa 100 ng of vector. The mix was incubated for 60 min at 50°C. Transformations were performed in Top10 competent bacteria (One Shot™ TOP10 Chemically Competent *E. coli*, Invitrogen) and selected using 100 μg/mL ampicillin or 50 μg/mL kanamycin sulphate (Euromedex). Agarose Gel purification and DNA plasmid extraction kits were purchased from Macherey-Nagel.

Dendra2_T69A_-LdcI and mGeosM-LdcI were both cloned in the pET-TEV vector containing an N-terminal 6xHis-tag, and a TEV cleavage site between Denda2_T69A_ or mGeosM and the LdcI gene. LdcI-Dendra2_T69A_ and LdcI-mGeosM were both cloned in the pET22b(+) vector with LdcI followed by either Dendra2_T69A_ or mGeosM containing an uncleavable C-terminal 6XHis-tag.

The anti-LdcI-Nb was obtained from the nanobody generation platform of the AFMB laboratory (Marseille, France). A llama (*Llama glama*) was immunized with the purified wild type LdcI. Lymphocytes were isolated from blood samples, a nanobody phage display library was generated, and enrichment of antigen-specific clones was performed using standard procedures (53, 54). Sequences of three positive clones were subcloned in the pHEN6 vector containing the pelB leader sequence from *Erwinia carotovora* for secretion into the periplasm, and a C-terminal 6xHis-tag (kindly provided by Dr. Aline Desmyter, AFMB Marseille). One of the clones was used for the present study to yield anti-LdcI-Nb.

Plasmids, primers and cloning strategy are summarised in Supplementary Table 4.

### Protein purification

LdcI-FP fusions and LdcI mutants were expressed in BL21(DE3) cells grown in LB medium supplemented with 100 μg/mL ampicillin or 50 μg/mL kanamycin sulphate. Protein expression was induced using 40 μM IPTG (Euromedex) and carried out overnight at 18°C. The LdcI-FP fusions and LdcI mutants were purified as previously described for wild-type LdcI (28, 29, 37), in a final buffer containing 25 mM Tris (pH 7.5), 0.3 M NaCl, 5% glycerol, 1 mM DTT and 0.1 mM PLP.

The anti-LdcI-Nb was expressed in *E. coli* WK6 cells following the protocol described by (53), and purified with immobilised metal affinity chromatography (IMAC, using a Ni-NTA column) followed by Size exclusion chromatography (SEC) using a a superdex 75 Increase 10/300GL column (GE-Healthcare) equilibrated with a buffer containing 25 mM Tris pH 7.4 and 0.3 M NaCl.

In order to characterise the LdcI complex with anti-LdcI-Nb, the two proteins were mixed at a 1:5 molar ratio and submitted to size exclusion chromatography as carried out for LdcI alone but without DTT in the buffer. The top of the peak was taken for subsequent ns-EM analysis.

### Ns-EM on LdcI-FP fusions, LdcI/anti-LdcI-Nb and LdcI mutants

FP fusion samples after gel filtration were diluted to a concentration of approximately 0.025 mg/mL. 3 μL was applied to the clean side of carbon on a carbon–mica interface and stained with 2% uranyl acetate (mGeosM-LdcI, LdcI-mGeosM, LdcI-Dendra2_T69A_, LdcI/anti-LdcI-Nb, LdcI mutants) or 2% sodium silicotungstate (Dendra2_T69A_-LdcI). Images were collected on a 120 kV Tecnai T12 microscope with an Orius 1000 camera (Gatan) or on a 200 kV Tecnai F20 electron microscope with either a OneView camera (Gatan) or a Ceta camera (Thermo Scientific). All images were collected with a defocus range of approximately −1.0 μm to −2.0 μm and with pixel sizes between 2.29 Å/pixel and 3.42 Å/pixel.

### Image processing – LdcI-FP and LdcI/anti-LdcI-Nb

36 micrographs of Dendra2_T69A_-LdcI with a pixel size of 2.73 Å/pixel, 368 micrographs of mGeosM-LdcI with a pixel size of 2.29 Å/pixel, 23 micrographs of LdcI-Dendra2_T69A_, 92 micrographs of LdcI-mGeosM with a pixel size of 3.42 Å/pixel, and 124 micrographs of LdcI/anti-LdcI-Nb with a pixel size of 2.82 Å/pixel were used for image analysis.

CTF estimation was performed with CTFFIND3 (55). Semi-automatic particle selection was carried out with BOXER (56), with box sizes of 98 × 98 pixels for Dendra2_T69A_-LdcI, 180 × 180 pixels for mGeosM-LdcI, 128 × 128 pixels for LdcI-Dendra2_T69A_ and LdcI-mGeosM, and 112 × 112 pixels for LdcI/anti-LdcI-Nb respectively. Particle extraction followed by several rounds of cleaning by 2D classification in RELION-1.4 (57), resulted in the following number of particles for each dataset: Dendra2_T69A_-LdcI = 7140, mGeosM-LdcI = 5514, LdcI-Dendra2_T69A_ = 832, LdcI-mGeosM = 12,211 and LdcI/anti-LdcI-Nb = 14,075.

For Dendra2_T69A_-LdcI, initial model generation was carried out in RELION-2.1 (58) without any symmetry applied. For mGeosM-LdcI, initial model generation was carried out in RELION-2.1 with either C1, C3 or D3 symmetry applied. The results of all three calculations being very similar, the model with applied D3 symmetry was selected. For LdcI-Dendra2_T69A_, LdcI-mGeosM and LdcI/anti-LdcI-Nb, the previously-determined LdcI decamer structure (PBD ID: 3N75) was filtered to 60 Å and used as an initial model for 3D refinement.

3D refinement was carried out for each dataset with applied C2 symmetry for Dendra2_T69A_-LdcI, D3 symmetry for mGeosM-LdcI, and D5 symmetry for LdcI-Dendra2_T69A_, LdcI-mGeosM and LdcI/anti-LdcI-Nb. Rigid body fitting of LdcI (PBD ID: 3N75), mEos2 (PDB ID: 3S05) and Dendra2 (PDB ID: 2VZX) crystal structures was then carried out in Chimera (59) for the five datasets.

### Cryo-EM on LdcI stacks (pH 5.7)

Wild-type LdcI was purified as previously described (10) from an *E. coli* strain impaired in the production of ppGpp (MG1655 Δ*relA*Δ*spoT*) in order to avoid any serendipitous ppGpp binding. Purified LdcI was diluted to a final concentration of approximately 0.25 mg/mL in a buffer containing 25 mM MES (pH 5.7), 0.3 M NaCl, 5% glycerol, 1 mM DTT and 0.1 mM PLP. 3 μL of the sample was applied to a glow-discharged R2/1 300 mesh holey carbon copper grid (Quantifoil Micro Tools GmbH) and plunge-frozen in liquid ethane using a Vitrobot Mark IV (FEI) operated at 100% humidity. Datasets were recorded at the European Synchrotron Radiation Facility (ESRF) in Grenoble, France (60), on a Titan Krios microscope (Thermo Scientific) equipped with a BioQuantum LS/967 energy filter (Gatan) and a K2 summit direct electron detector (Gatan) operated in counting mode. A total of 2564 movies of 30 frames were collected with a total exposure of 6 s, total dose of 29.3 e^-^/Å^2^ and a slit width of 20 eV for the energy filter. All movies were collected at a magnification of 130,000x, corresponding to a pixel size of 1.052 Å/pixel at the specimen level. A summary of cryo-EM data collection parameters can be found in Supplementary Table 1.

### Image Processing – LdcI stacks (pH 5.7)

Motion correction and dose-weighting of the recorded movies were performed using MotionCor2 (61). CTF parameters were determined on the aligned and dose-weighted sums using CTFFIND4 (62). After manual inspection of the dose-weighted sums, the best 558 (21.7%) micrographs were selected for further processing. LdcI stacks were manually picked using e2helixboxer in EMAN2 (63). A total of 15,165 LdcI-stack particles were extracted in RELION-3.0 (64) with an extract size of 320 pixels, resulting in boxes containing three LdcI decamers, and with the -- helix option with an outer diameter of 160 pixels and a helical rise of 77 pixels. After particle extraction, per-particle CTF correction was performed using Gctf (65). Extracted particles were subjected to 2D classification in RELION-3.0, resulting in a cleaned dataset containing 15,157 particles. Initial 3D refinement with imposed D5 symmetry was carried out in RELION-3.0, using an initial model generated by manually stacking three LdcI decamers (PDB ID: 3N75) in Chimera (59) and low-pass filtering the resulting LdcI-stack to 40 Å. The resulting 4.3 Å resolution 3D reconstruction, along with the cleaned particle stack, was subsequently imported into CryoSPARC (66). A final homogeneous 3D refinement in CryoSPARC, using a dynamic mask and imposing D5 symmetry, resulted in a map with a resolution of 3.28 Å based on the 0.143 gold-standard Fourier shell correlation (FSC) criterion (67). The final map was sharpened using a B-factor of −96 Å^2^. A local resolution estimation of the final 3D reconstruction was calculated in RELION-3.0. A summary of cryo-EM data collection parameters and image processing steps for LdcI stacks can be found in Supplementary Table 1 and Supplementary Figure 4, with local resolution and FSC curves shown in Supplementary Figure 5.

### Cryo-EM on LdcI decamers (pH 7.0)

Purified LdcI was dialysed into a buffer at pH 7.0 and diluted to a concentration of 0.25 mg/mL. 3 μL of diluted sample was applied to a glow-discharged (R2/1 300 mesh holey carbon copper grid (Quantifoil Micro Tools GmbH) and plunge-frozen in liquid ethane using a Vitrobot Mark IV (FEI) operated at 100% humidity. Images were recorded on a Glacios microscope (Thermo Scientific) equipped with a Falcon II direct electron detector (Thermo Scientific). A total of 2772 movies of 29 frames were collected with a total exposure of 1.5 s and a total dose of 45 e^−^/Å^2^. All movies were collected at a magnification of 116,086x, corresponding to a pixel size of 1.206 Å/pixel at the specimen level.

### Image Processing – LdcI decamers (pH 7.0)

Motion correction was carried out using patch motion correction in CryoSPARC, discarding the first two frames. Initial CTF estimation was then carried out on summed frames using CTFFIND4. A subset of 600 micrographs were subjected to automatic picking using the blob picker in CryoSPARC, resulting in ~238,000 picked particles. Particles were then extracted with a box size of 256 × 256 and subjected to 2D classification. Particles from the best classes showing clear secondary structural features for LdcI were selected for homogeneous refinement (with applied D5 symmetry) against EMD-3204 low-pass filtered to 30 Å, resulting in a reconstruction with a resolution of 4.2 Å (FSC = 0.143). This reconstruction was then used to create templates for picking the entire dataset using the template picker in CryoSPARC, after filtering to 12 Å. ~796,000 particles were extracted and subjected to 2D classification, and the best ~428,000 particles were subjected to heterogeneous refinement with applied D5 symmetry against the 4.2 Å map, resulting in one higher-resolution class corresponding to ~229,000 particles. These particles were subjected to homogeneous refinement with applied D5 symmetry, resulting in a map with a resolution of 2.76 Å (FSC = 0.143) which was then sharpened with a B-factor of −173 Å^2^. A summary of cryo-EM data collection parameters and image processing steps for LdcI decamers can be found in Supplementary Table 1 and Supplementary Figure 4, with local resolution and FSC curves shown in Supplementary Figure 5.

### Fitting of structures and refinement

For fitting of atomic models in the 3D reconstructions of LdcI at pH 5.7 (stack) or pH 7.0 (decamer), two (for the stack) or one copy (for the decamer) of the LdcI X-ray crystal structure (PDB ID: 3N75) were first rigid-body fitted in the corresponding 3D reconstructions using Chimera. Refinement was performed using the Phenix software package (68) and was identical for both 3D reconstructions. A first round of real space refinement was carried out with enabled rigid-body, global minimization, local grid search and ADP refinement parameters, and imposing rotamer, Ramachandran, NCS and reference model (PDB ID: 3N75) restraints. A final round of real space refinement was then performed using the same settings, but without rigid body refinement and without applying reference restrains, setting the ‘nonbonded_weight’ parameter to 4000 and disabling ‘local_grid_search’. A summary of refinement and model validation statistics can be found in Supplementary Table 1.

### pH shift experiment

Stationary phase cultures, which were grown overnight from single colonies in LB medium, were diluted to OD600 ~ 0.01 and re-grown at 37°C within approximately 1h 45 min in fresh LB medium to an OD600 of 0.1. From this culture, 14 mL were transferred to 15 mL falcon tubes and pelleted by centrifugation at RT for 5 min. The supernatant was decanted, whereby systematically around 200 μL LB remained in the falcon tube. The pellet of bacteria was resuspended and afterwards, LB-4.6 containing 30 mM L-lysine was added to the cells up to 14 mL. To prepare LB-4.6 medium, LB powder (Sigma-Aldrich) was completely dissolved in distilled water by stirring for 30 min. The pH of 4.6 was then adjusted using HCl. After autoclaving, sterile filtered L-lysine was added to LB-4.6. For this, L-lysine was dissolved in an aliquot of LB-4.6, sterile filtered and mixed with the remaining LB-4.6. 30 mM L-lysine were used. In order to grow the culture under oxygen-limiting conditions, the lid of the falcon tube was closed and the tubes were placed at 37°C on a shaker (150 rpm). After defined time-points, aliquots were taken for OD600 measurement, pH measurement, SDS-PAGE or immunofluorescence. For each time-point, a new tube was opened and not reused further.

### Western Blotting

SDS-PAGE was performed with a Biorad electrophoresis chamber using standard 12% reducing SDS-PAGE gels. Proteins were transferred to a nitrocellulose membrane (Biorad) using a Trans-Blot Turbo Transfer System (Biorad). The membrane was blocked for 1h using 5% BSA in TBS supplemented with 0.1% Tween (TBS-T). Afterwards, the membrane was incubated for 1h with an anti-LdcI antibody (Qalam-Antibodies) in BSA/TBS-T (1:5000). The membrane was subsequently washed 3 × 10 min in TBS-T and incubated for 1 h with HRP-coupled anti-rabbit antibody (1:10000 in BSA/TBS-T). Finally the membrane was washed 3 × 10 min in TBS-T. The membrane was rinsed once with TBS, prior to detection of antibody-labelled proteins using ECL reagent (GE Heathcare). Incubation and detection of antibody was performed at 25 °C.

### Nanobody labelling

For the labelling reaction, 50 μL nanobody (i.e. 200 μg) was pipetted into a 1.5 mL tube, placed on ice and supplemented with 5.5 μL of 1 M bicarbonate buffer at pH 8.3. 100 μg of Alexa-647 or Alexa-488 NHS ester dye (Life Technologies, A37573 and A20000) were dissolved in 10 μL DMSO to final concentration of 10 mg/mL. 5 μl (i.e. 40 nmol) dye in DMSO was added to the protein and incubated for 1 h at room temperature in a shaking block, covered with an aluminium foil to protect the dye from the light. Excess dye was removed by iterative buffer exchange using a 3K spin column (Amicon-Ultra-4 Centrifugal Filters Ultracell 3K, Millipore UFC800396) to PBS. The degree of labelling was inferred from measuring the OD.

### Cell preparation for immunofluorescence staining of *E. coli* cells with nanobodies

For immunofluorescence staining, cells were subjected to a pH shift in order to induce LdcI expression, as described in the “pH shift experiment” section above. 90 minutes after the pH shift, the OD600 of the cell culture was measured, and the volume of cells corresponding to OD = 4, with OD = 1 corresponding to about 8×10^8^ cells per mL, were collected by centrifugation. After removal of LB by pipetting, cells were resuspended in 2 mL of 4 % FA in PBS (made from 16% Formaldehyde solution, Methanol-free from Thermo Scientific). Falcons were placed on a rotor for constant agitation for 45 min at room temperature. After fixation, cells were collected by centrifugation and the solution was removed by pipetting. Cells were then resuspended in 14 mL PBS (Gibco, Thermo Scientific) to remove and dilute the fixative. Cells were permeabilised for 10 min using 2 mL 0.1 % Triton X-100 in PBS and subsequently washed three times with 10 mL PBS. Finally, cells were transferred to 1.5 mL tubes, centrifuged and resuspended in 200 μL 1% BSA/PBS (BSA/PBS solution was dissolved for 30 min and sterile-filtered to avoid clumps). After 30 min of incubation, 0.5 μg anti-LdcI-Nb, labelled with the dedicated dye, was added to the 200 μL bacteria-BSA/PBS suspension. Cells were incubated with the labelled anti-LdcI-Nb for 16 h at 4°C. The next day, cells were washed three times with 1 mL PBS, centrifuged to remove antibody solution and resuspended in 250 μl PBS. When needed, Hoechst 33342 (Sigma-Aldrich) was added to a final concentration of about 100 ng/mL.

### Wide-field imaging and flow cytometry

For epifluorescence imaging, 2 μL of cells were placed between a glass slide and a coverslip, which have been carefully pressed together, and observed using an inverted IX81 microscope, with a UPLFLN 100X oil immersion objective (N.A. 1.3) (Olympus), using the appropriate specific excitation and emission filters for AF488 (GFP-3035B set, Semrock) and DAPI (DAPI-5060B set, Semrock). Acquisitions were performed with Volocity software (QuorumTechnologies™) with a sCMOS 2048×2048 camera (Hamamatsu ORCA Flash 4, 16 bits/pixel) achieving a final magnification of 64 nm per pixel. For flow cytometry, 50 μL of cells suspensions were injected in a MACSQuant VYB flow cytometer (Miltenyi Biotech, Bergish Gladbach, Germany) using the 488 nm excitation and 525 (50) nm emission channel (B1). AF488 positive populations were estimated after forward scatter (FSC) and side scatter (SSC) gating on the cells. Data were further processed with MACSQuantify software (Miltenyi Biotech).

### STORM imaging

For Single Molecule Localization Microscopy, cells were transferred to a glucose buffer containing 50 mM NaCl, 150 mM Tris (pH 8.0), 10% Glucose, 100 mM MEA (Mercaptoethylamine) and 1x Glox. Glox was prepared as a 10x stock and contained 1 μM catalase and 2.3 μM glucoseoxidase. Menzel glass slides (Thermo Scientific) and precision coverslips (1.5H from ThorLabs) were cleaned for 30 min using an UV-ozone cleaning device (HELIOS‐500, UVOTECH Systems). 2 μL of immuno-labelled cells were placed onto a glass slide and covered with the coverslip, then cells were carefully spread by pressing glass slides firmly together and the sides were sealed with transparent nail polish to avoid evaporation. Mounted samples were imaged on a homemade SMLM set up based on an IX81 microscope (Olympus). STORM was performed by focusing a 643 nm excitation laser beam (Toptica Diode laser) to the back focal plane of an oil immersion UAPON100X (N.A. 1.49) objective. The intensity of the laser was tuned using an Acousto-Optical Tunable Filter (OATF, Quanta Tech). Acquisition was obtained with a 16 bits/pixel Evolve 512 EMCCD (Photometrics) using Metamorph (Molecular Devices), for a final pixel size of 123 nm. 3D-STORM based on point-spread-function astigmatism (69) was performed using a cylindrical lens (LJ1516L1-A, Thorlabs) placed in the detection light path. STORM datasets consisting of about 30,000 frames, using a 643 nm laser power density of 3 kW/cm² and a 405 nm laser power density of up to 1 W/cm^2^ and a frametime of 50 ms, were acquired with an EMCCD gain set to 200. 3D point spread function calibration was achieved using tetraspec beads. Finally, data were processed with the Thunderstorm plugin (70) in ImageJ (71). 3D-images were rendered with Visp (72) using a minimum neighbour density threshold of 20 to 28. A total of about 25,000 localisations with a median localisation precision of ~17 nm were recorded per cell. A quantitative evaluation of molecular copy number from STORM data is notoriously difficult due to pronounced and potentially environmentally-dependent blinking of AF647. However, assuming an average of 3 blinking events per molecule and 100% efficient labelling, one can estimate the number of LdcI decamers in the range 500-1000 per cell, which would agree very well with earlier independent estimations (6, 31). Therefore, although the LdcI labelling efficiency *in cellulo* was not precisely determined in this study, it is likely to be quite high, which is also consistent with our *in vitro* structural data on the LdcI-nanobody complex.

### Biolayer interferometry measurements

For BLI binding studies, RavA with a biotinylated C-terminal AviTag was expressed and purified as previously described (30). BLI experiments were performed in 1x HBS pH at 7.0 (25 mM HEPES, 300 mM NaCl, 10 mM MgCl_2_, 10% glycerol) supplemented with 1x kinetics buffer (0.1% w/v BSA, 0.02% v/v Tween-20), 1mM ADP, 1mM DTT and 0.1 mM PLP. Experiments were performed using the BLItz System instrument (FortéBio), operated at room temperature. Before the start of each BLI experiment, RavA-AviTag was incubated with 1 mM ADP for 10 min. Streptavidin-coated biosensors (FortéBio) were functionalised with biotinylated RavA-AviTag, then quenched with 10 μg/mL biocytin. Experiments with the wild-type LdcI are from (30). For C-terminal LdcI-FP fusions, pins were dipped in wells containing a range of LdcI-FP concentrations from 0 to 1000 nM, with no binding signal recorded at any concentration of LdcI-FP.

## Data and code availability

Cryo-EM maps, along with the corresponding fitted atomic structures, have been submitted to the EMDB and PDB with accession codes EMD-10850 and PDB-6YN6 for LdcI stacks (pH 5.7), and EMD-10849 and PDB-6YN5 for LdcI decamers (pH 7.0).

## Acknowledgements

We thank Guy Schoehn for establishing and managing the cryo-electron microscopy platform and for providing training and support, and Rose-Laure Revel-Goyet and Françoise Lacroix for the support and access to the M4D Cell imaging Platform. We are grateful to Aymeric Peuch for help with the usage of the EM computing cluster and Daniel Thédie, Kévin Floc’h and Joel Beaudoin for discussions. We thank Alain Roussel and Aline Desmyter for the nanobody production, and Julien Perard for help with the cloning of the Dendra2_T69A_-LdcI construct. We acknowledge the European Synchrotron Radiation Facility for provision of beam time on CM01. This work was funded by the European Union’s Horizon 2020 research and innovation programme under grant agreement No 647784 to IG. The nanobody generation platform of the AFMB laboratory (Marseille, France) was supported by the French Infrastructure for Integrated Structural Biology (FRISBI) ANR-10-INSB-05-01. For electron and fluorescence microscopy studies, this work used the platforms of the Grenoble Instruct centre (ISBG; UMS 3518 CNRS-CEA-UJF-EMBL) with support from FRISBI (ANR-10-INSB-05-02) and GRAL (ANR-10-LABX-49-01) within the Grenoble Partnership for Structural Biology (PSB). The electron microscope facility (Glacios electron microscope) is supported by the Rhône-Alpes Region (CIBLE and FEDER), the FRM, the CNRS, the University of Grenoble and the GIS-IBISA. JF was supported by a long-term EMBO fellowship (ALTF441-2017) and a Marie Skłodowska-Curie actions Individual Fellowship (789385, RespViRALI).

## Author Contributions

C.L., M.B., V.A., A.F. and K.H cloned constructs, M.J., A.F. and K.H. purified proteins. M.J. and I.G. performed ns-EM imaging and analysis. A.D., G.E. and M.B.-V. performed cryo-EM imaging. J.F. and A.D. performed cryo-EM analysis. M.J. and J.F. built models resulting from EM maps. M.J., J.F. and I.G. analysed structures and interpreted data. C.L. and M.B. performed optical imaging of overexpression constructs with input of V.A., J.-P.K. and D.B. C.L. performed nanobody characterisation for optical imaging with input from J.-P.K. C.L. analysed endogenous expression and performed optical imaging of endogenous LdcI with input of V.A., J.-P.K. and D.B. C.L. and D.B. analysed STORM Images. M.J., C.L., J.F. and I.G. analysed the data and prepared the figures and tables. C.L. and D.B. contributed to the design of the optical imaging part of the project together with I.G. I.G. designed, supervised and funded the overall study. M.J., J.F. and I.G. wrote the manuscript with significant input from C.L. and contributions from all of the authors.

---

**Supplementary Figure 1.**
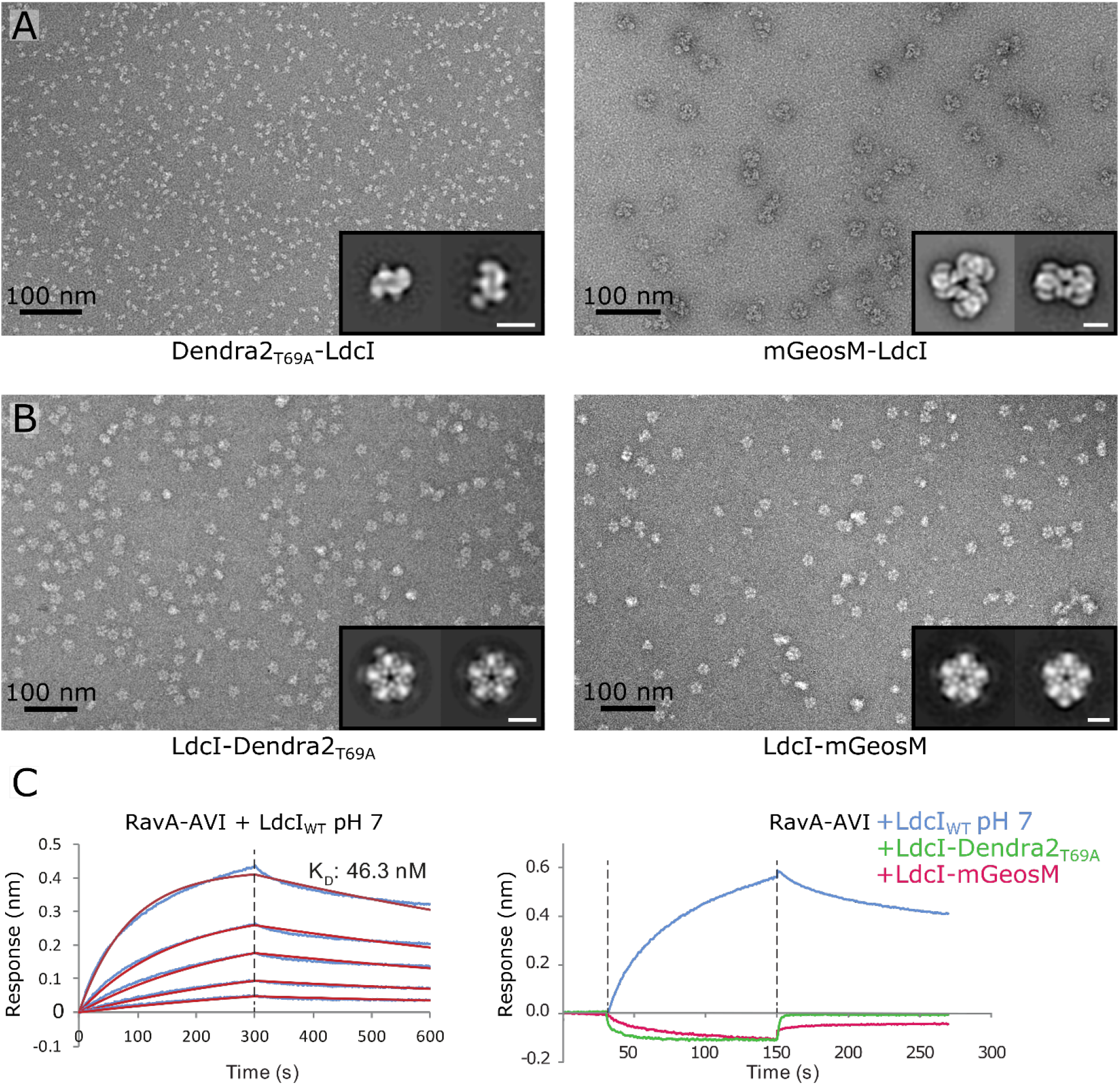
Characterisation of Dendra2_T69A_-LdcI and mGeosM-LdcI fusion constructs by negative stain EM and BLI. **A)** Ns-EM micrographs of N-terminal FP-LdcI fusions Dendra2_T69A_-LdcI (left) and mGeosM-LdcI (right), with 2D class averages inset (scale bar = 100 Å). Dendra2_T69A_-LdcI only forms dimers, compared to mGeosM-LdcI which forms large non-native dodecamers. **B)** Ns-EM micrographs of C-terminal LdcI-FP fusions LdcI-Dendra2_T69A_ (left) and LdcI-mGeosM (right), with 2D class averages inset (scale bar = 100 Å). Both fusions form decamers, with flexible-linked fluorophores visible as weak densities around the outside of the decameric ring in the 2D class averages. **C)** BLI binding curves of wild-type LdcI (left, [LdcI_WT_] = 500, 250, 125, 62.5, and 31.25 nM) and LdcI_WT_ (right, 500 nM), LdcI-Dendra2T69A-LdcI (right, 500 nM) and LdcI-mGeosM-LdcI (right, 500 nM) against RavA-AVI. In contrast to wild-type LdcI, which binds RavA-AVI with a 46.3 nM affinity, there is no measurable interaction between either of the C-terminal LdcI-FP fusions and RavA-AVI.

**Supplementary Figure 2.**
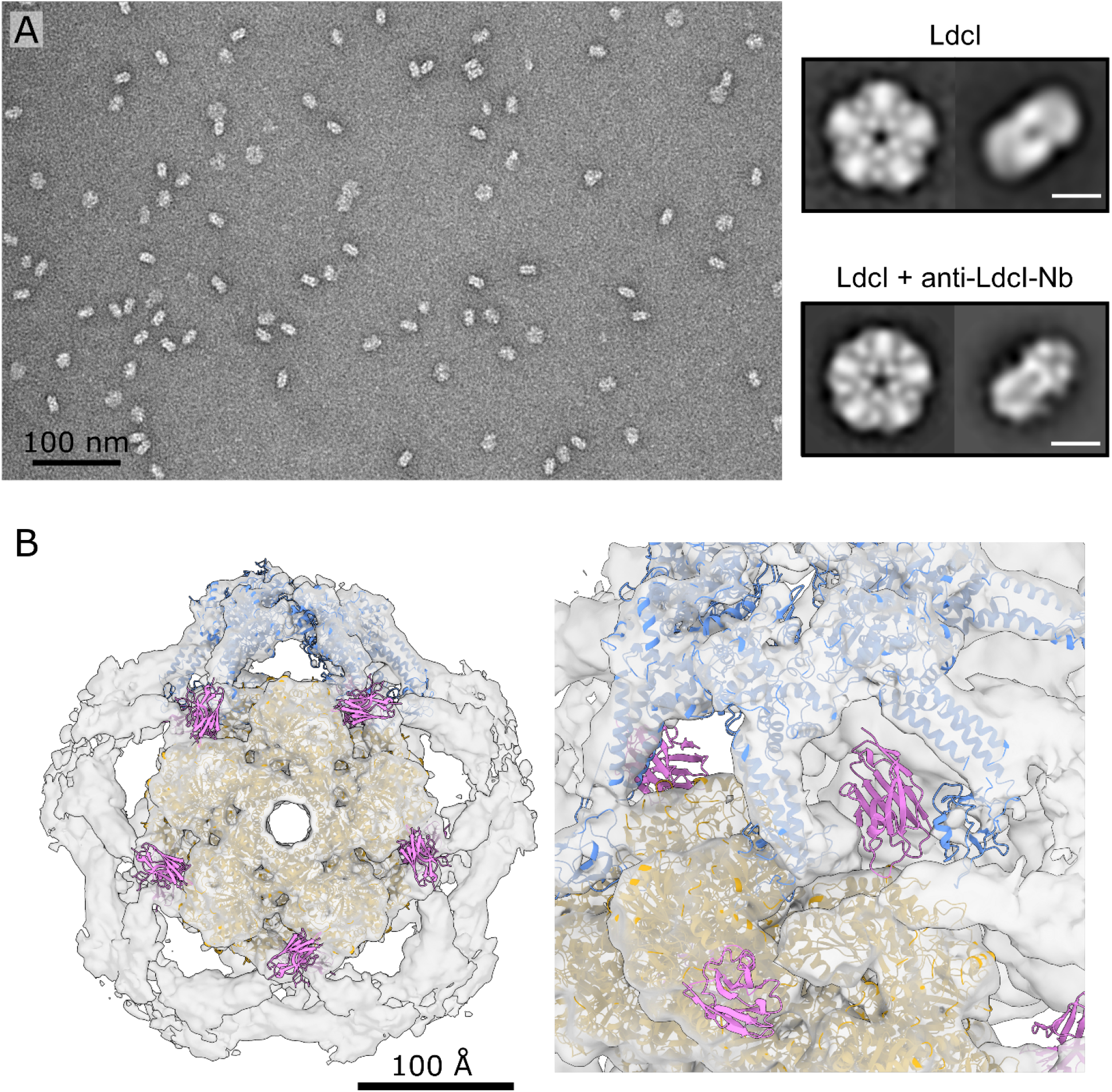
LdcI interacts with anti-LdcI-Nb and RavA at distinct sites. **A)** Ns-EM micrograph of LdcI + anti-LdcI-Nb (left), with 2D class averages (right) of LdcI alone and LdcI + anti-LdcI-Nb (scale bar = 100 Å). **B)***In silico* model of anti-LdcI-Nb (represented by the anti-lysozyme nanobody crystal structure, PDB ID: 1MEL) superimposed over the cryo-EM map of the LdcI-RavA complex (Jessop et al., 2020) (EMDB ID: 4469, PDB ID: 6Q7L). The nanobody (coloured magenta) would occupy a distinct location to the RavA binding site in the LdcI-RavA cage-like complex (coloured gold and blue).

**Supplementary Figure 3.**
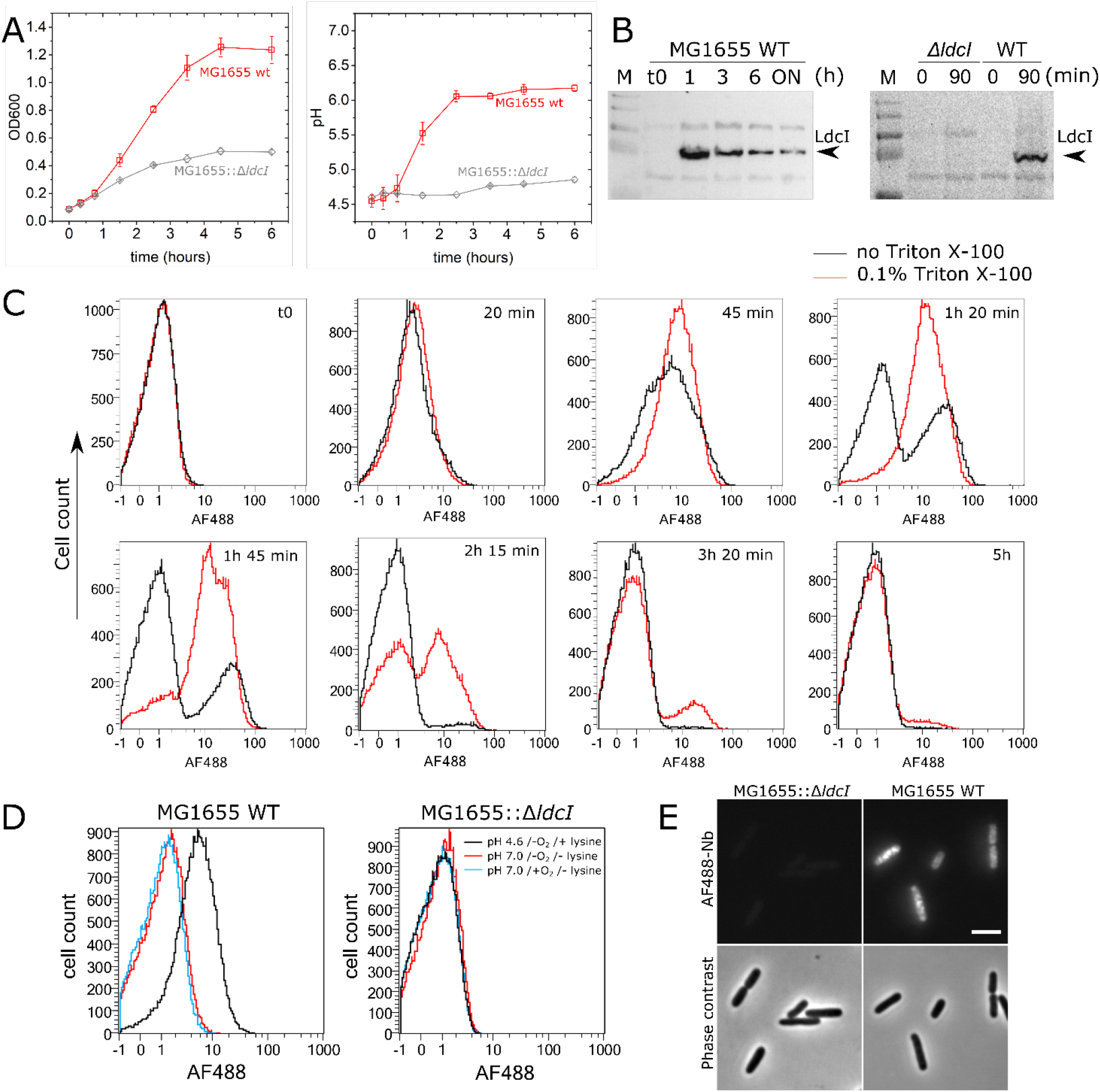
Immunolabelling of endogenous LdcI in *E. coli* upon acid stress. Characterisation of LdcI expression using anti-LdcI-Nb. **A)** Left – growth curve of wild-type (red) and LdcI knock-out (grey) cells after exposure to acid stress (pH 4.6), showing compromised growth in LdcI KO cells. Right – measured pH of the growth medium for wild-type and LdcI KO cells after exposure to acid stress (pH 4.6). Wild-type cells are able to buffer the medium under acid stress, whereas LdcI knock-out cells are not. Each data point represents the average of 3 independent measurements taken from 2 biological replicates, error bars correspond to the calculated standard deviation. **B)** Left – western blot showing LdcI expression after exposure to acid stress after 0, 1, 3 and 6 hours, and overnight (ON). LdcI expression is maximal soon after exposure to acid stress, tailing off after three hours. Right – western blot against LdcI in *ΔldcI* and WT MG1655 cells, 0 minutes and 90 minutes after exposure to acid stress, showing a lack of LdcI expression in *ΔldcI* cells and a lack of baseline expression before exposure to acid stress in WT cells. M = molecular weight marker. **C)** Flow cytometry measurements carried out after exposure of wild-type cells to acid stress show that LdcI is maximally expressed one to two hours after exposure to acid stress, before tailing off. **D)** Flow cytometry measurements of wild-type and LdcI knock-out cells labelled with anti-LdcI-Nb-AF488 after 90 minutes of growth under three conditions – pH 4.6 in the absence of oxygen and lysine, neutral pH in the absence of oxygen and lysine, and neutral pH in the presence of oxygen but absence of lysine. No fluorescence is seen above baseline for the LdcI knock-out cells under any condition. For wild-type cells, fluorescence is only seen under acid stress conditions. **E**) Wide-field fluorescence microscopy of LdcI knock-out MG1655 cells (left) and wild-type MG1655 cells (right), 90 minutes after exposure to acid stress. The staining of anti-LdcI-Nb-AF488 is specific, shown by the lack of background in the knock-out strain. Scale bar = 2μm.

**Supplementary Figure 4.**
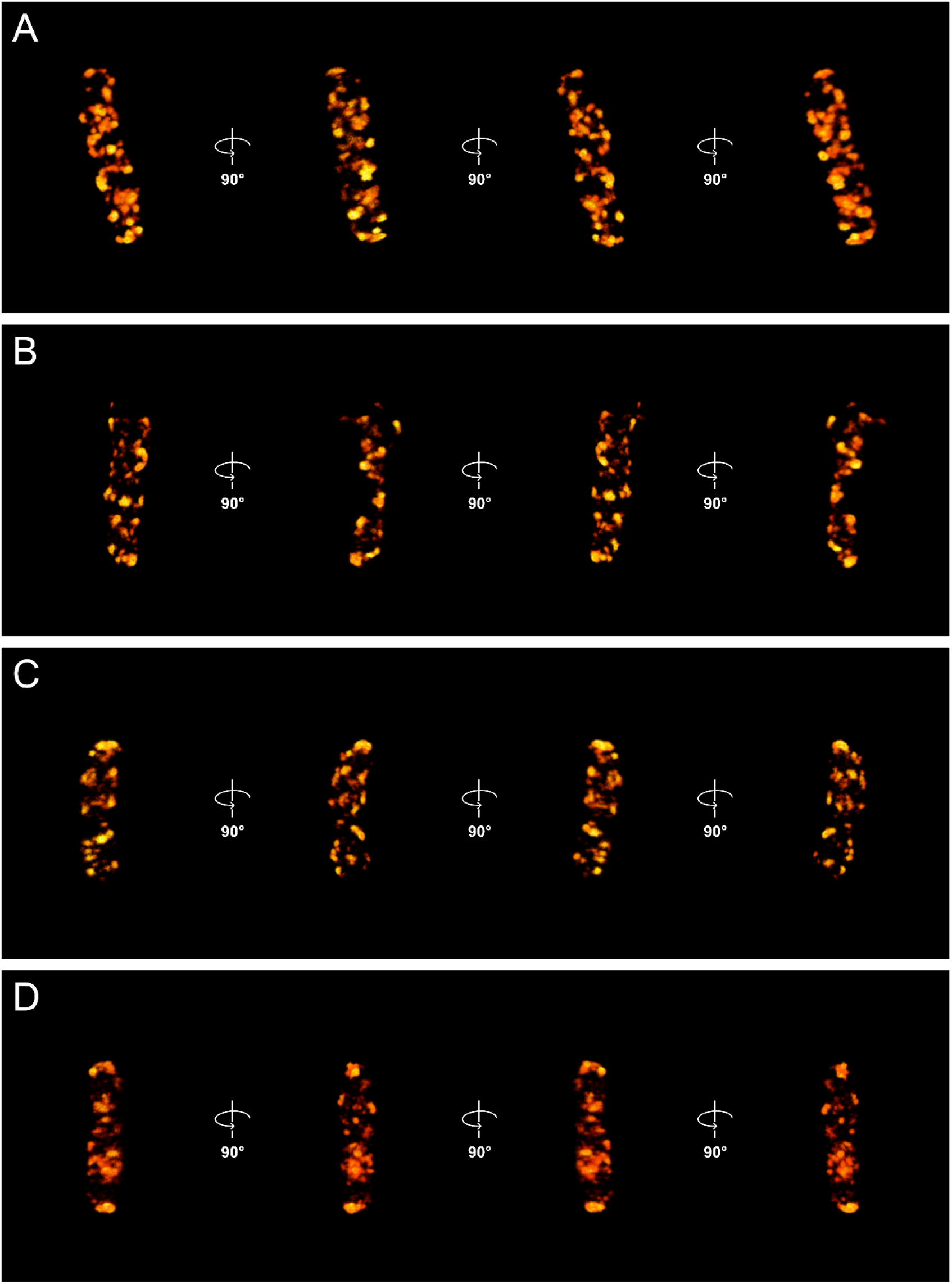
Galleries of 3D-STORM images of the four individual *E. coli* cells shown in Figure 3. Different orientations of each cell related by 90° rotations around the long axis of the bacterium are shown in order to facilitate visualisation of the patchy distribution of LdcI.

**Supplementary Figure 5.**
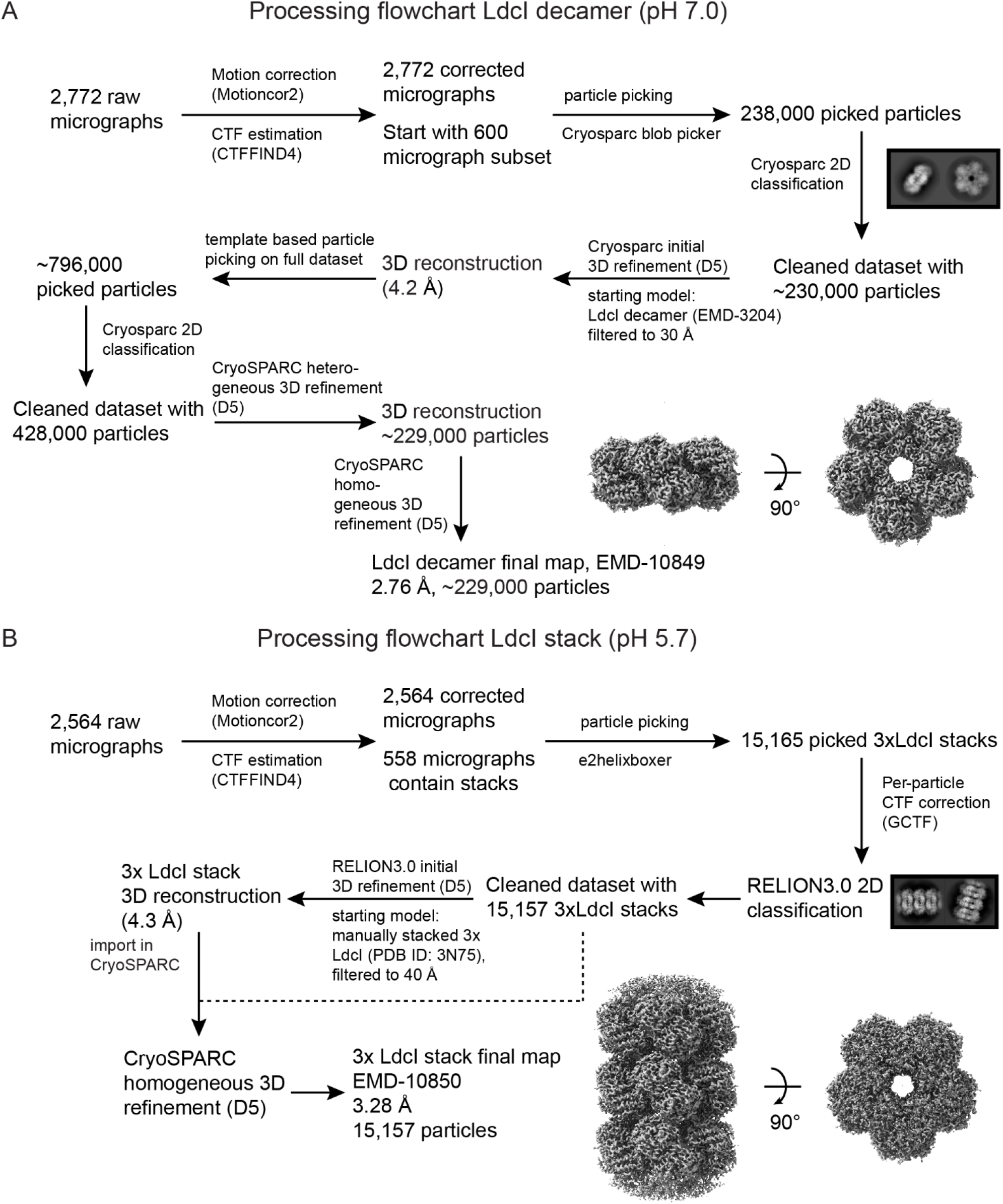
Cryo-EM processing pipelines. **A)** Cryo-EM processing pipeline for LdcI decamers at pH 7.0. Unless otherwise indicated, processing steps were carried out in CryoSPARC. **B)** Cryo-EM processing pipeline for LdcI stacks at pH 5.7. Software packages used at each step are indicated. For a summary of data collection and processing parameters, and refinement and validation statistics, see Supplementary Table 1.

**Supplementary Figure 6.**
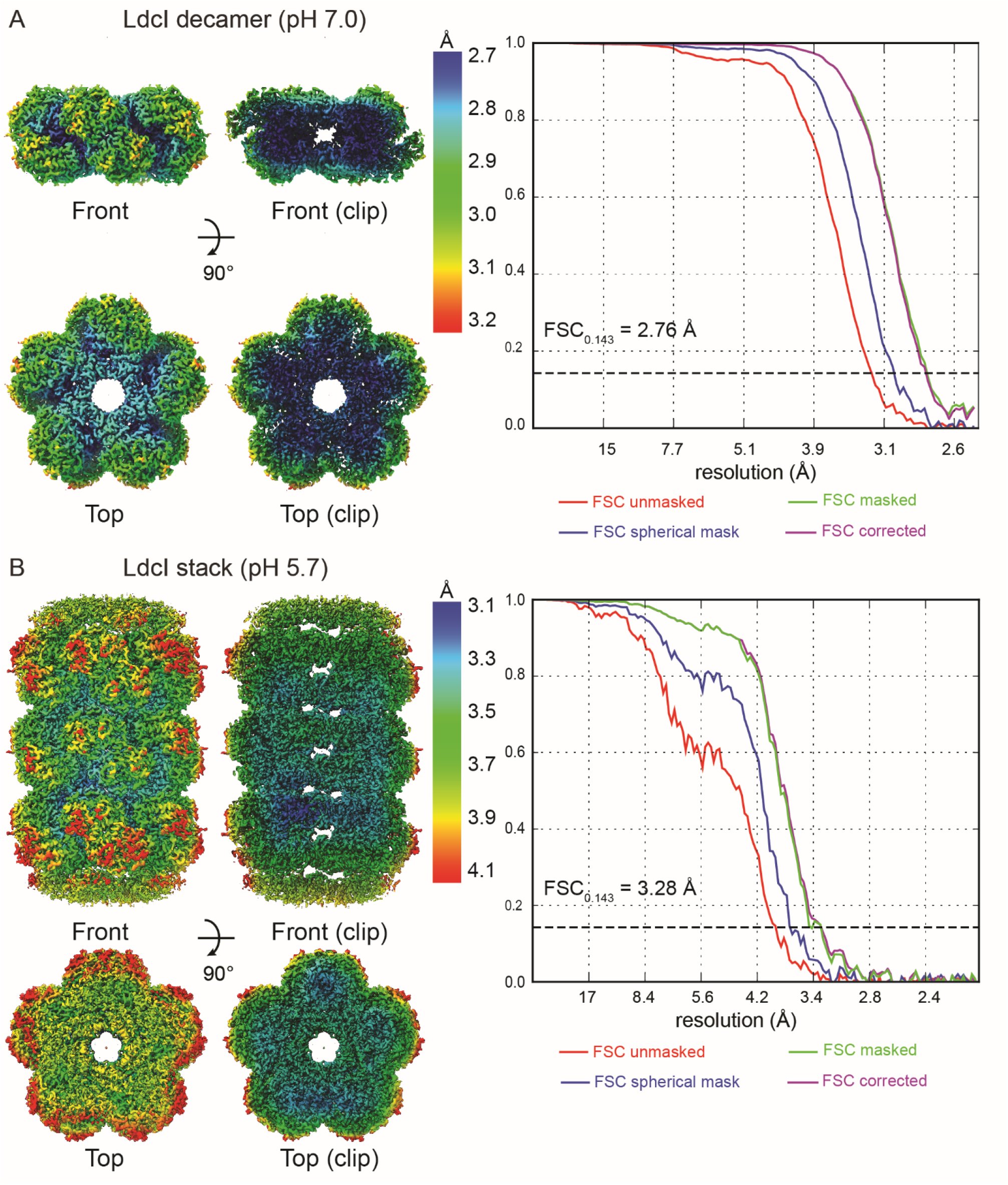
Assessment of the resolution of the cryo-EM maps. 3D reconstructions of **A)** the LdcI decamer at pH 7.0 **B)** the LdcI stack at pH 5.7, coloured according to the local resolution. Gold-standard FSC curves with the estimated resolution at FSC = 0.143 are shown on the right.

**Supplementary Figure 8.**
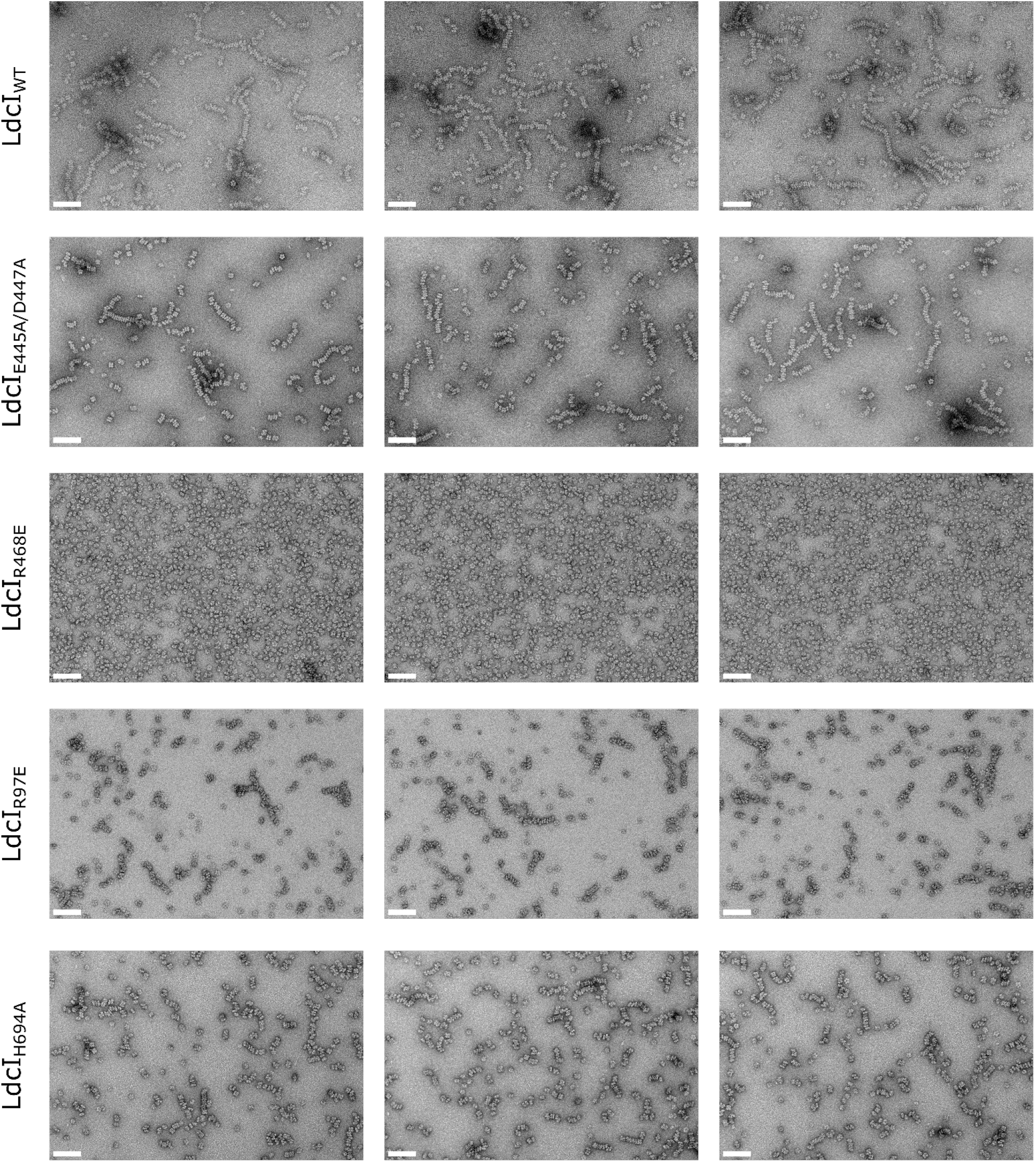
Mutational analysis of the predicted molecular determinants of LdcI polymerisation under acid stress conditions. Full uncropped ns-EM micrographs of wild type and mutant LdcI at pH 5.7, scale bars = 100 nm. Three representative micrographs are shown for each construct. Micrographs for LdcI_R97E/R468E_, LdcI_E445A/D447A/R468E_, and LdcI_H694N_ were also collected but are not shown here. LdcI_E445A/D447A/R468E_ and LdcI_R97E/R468E_ behaved like LdcI_R468E_ and remained entirely decameric at low pH, while LdcI_H694N_ displayed similar behaviour to LdcI_H694A_.

**Supplementary Table 1.** Cryo-EM data collection, refinement and validation statistics.

|  | Ldcl stacks<br>(pH 5.7) | Ldcl decamers<br>(pH 7.0) |
| --- | --- | --- |
|  | EMDB 10850<br>PDB 6YN6 | EMDB 10849<br>PDB 6YN5 |
| **Data collection and processing** |  |  |
| Magnification | 130,000 | 116.086 |
| Voltage (kV) | 300 | 200 |
| Electron exposure (e-/Å<sup>2</sup>) | 29.3 | 45.0 |
| Defocus range (µm) | 1.0 – 2.5 | 1.0 – 3.0 |
| Pixel size (Å) | 1.052 | 1.206 |
| Symmetry imposed | D5 | D5 |
| Initial particle images (no.) | 15,165 | ~796,000 |
| Final particle images (no.) | 15,157 | ~229,000 |
| Map resolution (Å) | 3.3 | 2.8 |
| FSC threshold | 0.143 | 0.143 |
| Map resolution range (Å) | **69.0-3.3** | **51.2-2.8** |
| **Refinement** |  |  |
| Initial model used (PDB code) | 3N75 | 3N75 |
| Model resolution (Å) (masked) | 3.7 | 2.9 |
| FSC threshold | 0.5 | 0.5 |
| Model resolution range (Å) | ∞-3.7 | ∞-2.9 |
| Map sharpening *B* factor (Å<sup>2</sup>) | -96 Å<sup>-2</sup> | -173 Å<sup>-2</sup> |
| Model composition |  |  |
| Non-hydrogen atoms | 113,760 | 57,074 |
| Protein residues | 14,220 | 7,110 |
| Ligands | 0 | 0 |
| *B* factors (Å<sup>-2</sup>) |  |  |
| Protein | 91.22 | 36.51 |
| Ligand | - | - |
| R.m.s. deviations |  |  |
| Bond lengths (Å) | 0.009 | 0.012 |
| Bond angles (°) | 1.131 | 1.268 |
| Validation |  |  |
| MolProbity score | 1.59 | 1.28 |
| Clashscore | 4.78 | 2.83 |
| Poor rotamers (%) | 0.49 | 0.19 |
| Ramachandran plot |  |  |
| Favored (%) | 95.04 | 96.74 |
| Allowed (%) | 4.96 | 3.26 |
| Outliers (%) | 0.00 | 0.00 |

**Supplementary Table 2.** Root-mean-square-deviation (RMSD) between extracted LdcI monomer structures at pH 8.5 (PDB ID: 3N75), 7.0 and 5.7.

|  | Ldcl decamer, pH 8.5 | Ldcl decamer, pH 7.0 | Ldcl decamer, pH 5.7 |
| --- | --- | --- | --- |
| Ldcl decamer, pH 8.5 | 0 | 0.24 Å (670 aligned atoms) | 0.61 Å (610 aligned atoms) |
| Ldcl decamer, pH 7.0 | 0.24 Å (670 aligned atoms) | 0 | 0.56 Å (604 aligned atoms) |
| Ldcl decamer, pH 5.7 | 0.61 Å (610 aligned atoms) | 0.56 Å (604 aligned atoms) | 0 |

**Supplementary Table 3.** Root-mean-square-deviation (RMSD) between extracted LdcI dimer structures at pH 8.5 (PDB ID: 3N75), 7.0 and 5.7.

|  | Ldcl decamer, pH 8.5 | Ldcl decamer, pH 7.0 | Ldcl decamer, pH 5.7 |
| --- | --- | --- | --- |
| Ldcl decamer, pH 8.5 | 0 | 0.30 Å (1328 aligned atoms) | 1.51 Å (1281 aligned atoms) |
| Ldcl decamer, pH 7.0 | 0.30 Å (1328 aligned atoms) | 0 | 1.41 Å (1287 aligned atoms) |
| Ldcl decamer, pH 5.7 | 1.51 Å (1281 aligned atoms) | 1.41 Å (1287 aligned atoms) | 0 |

**Supplementary Table 4, A.**
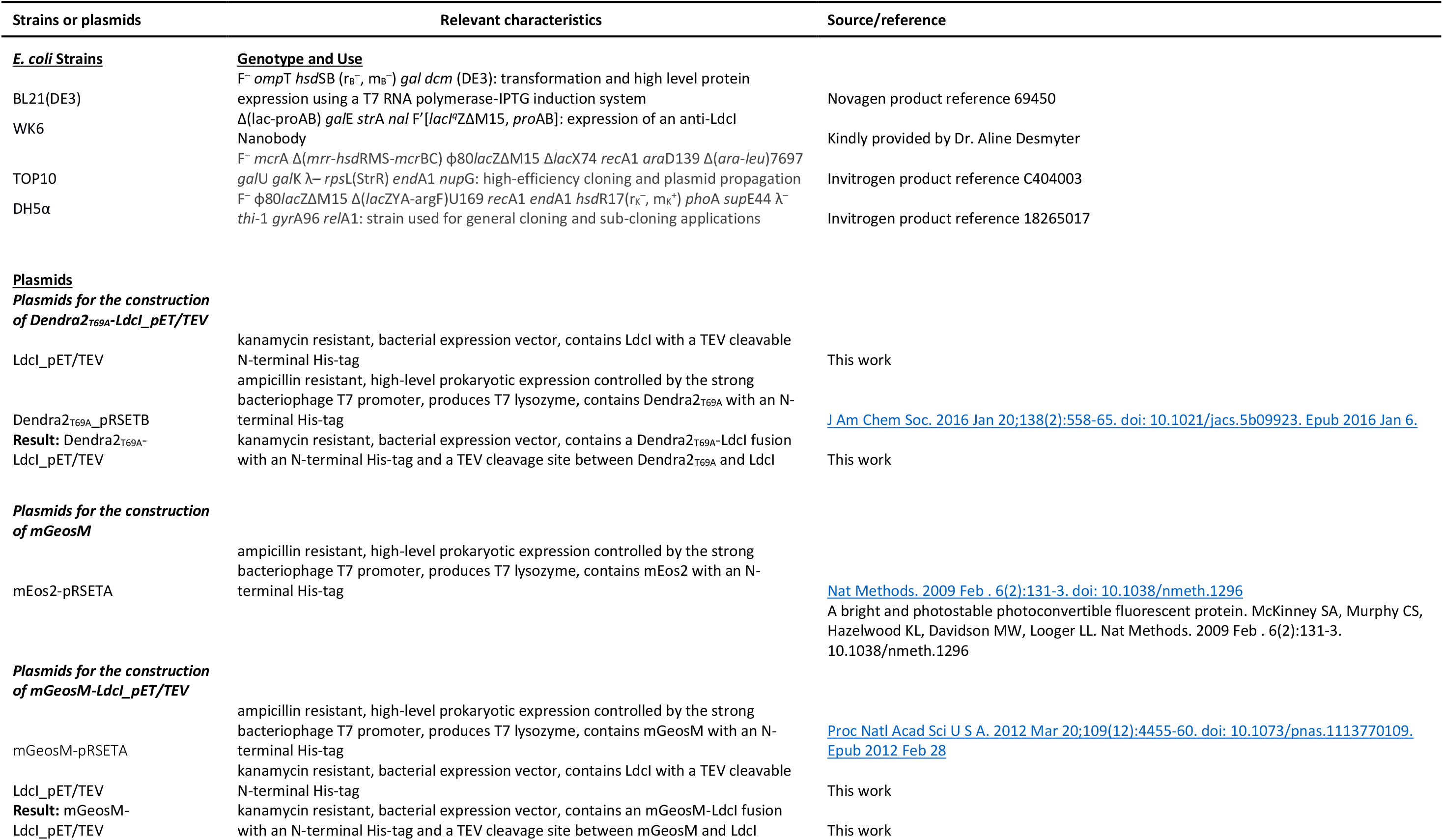
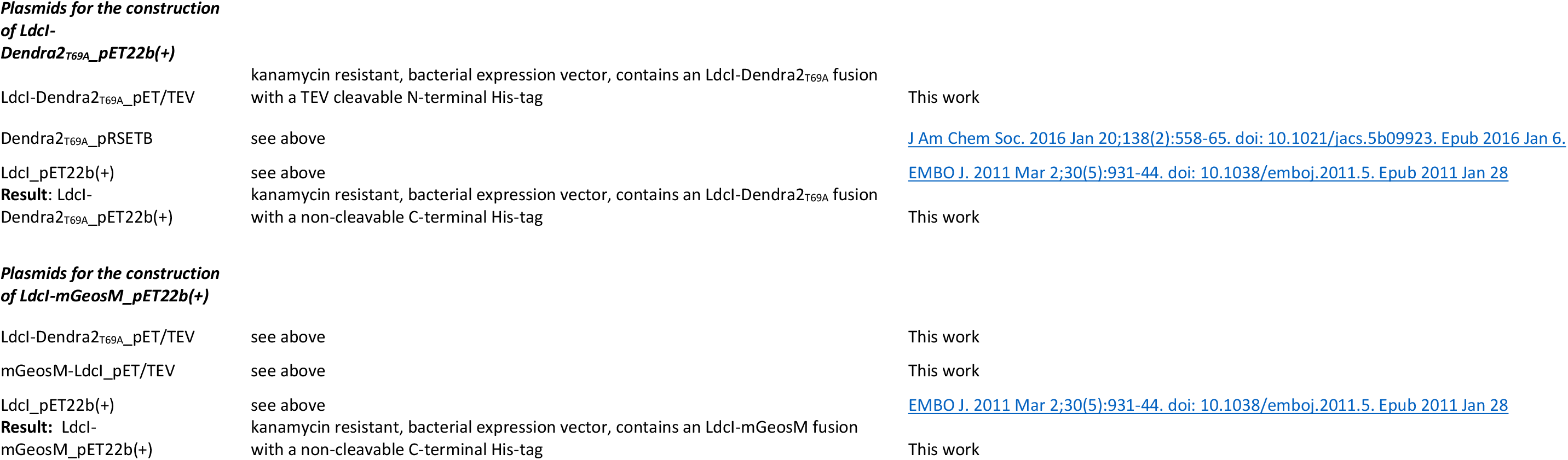

**Supplementary Table 4, B.**
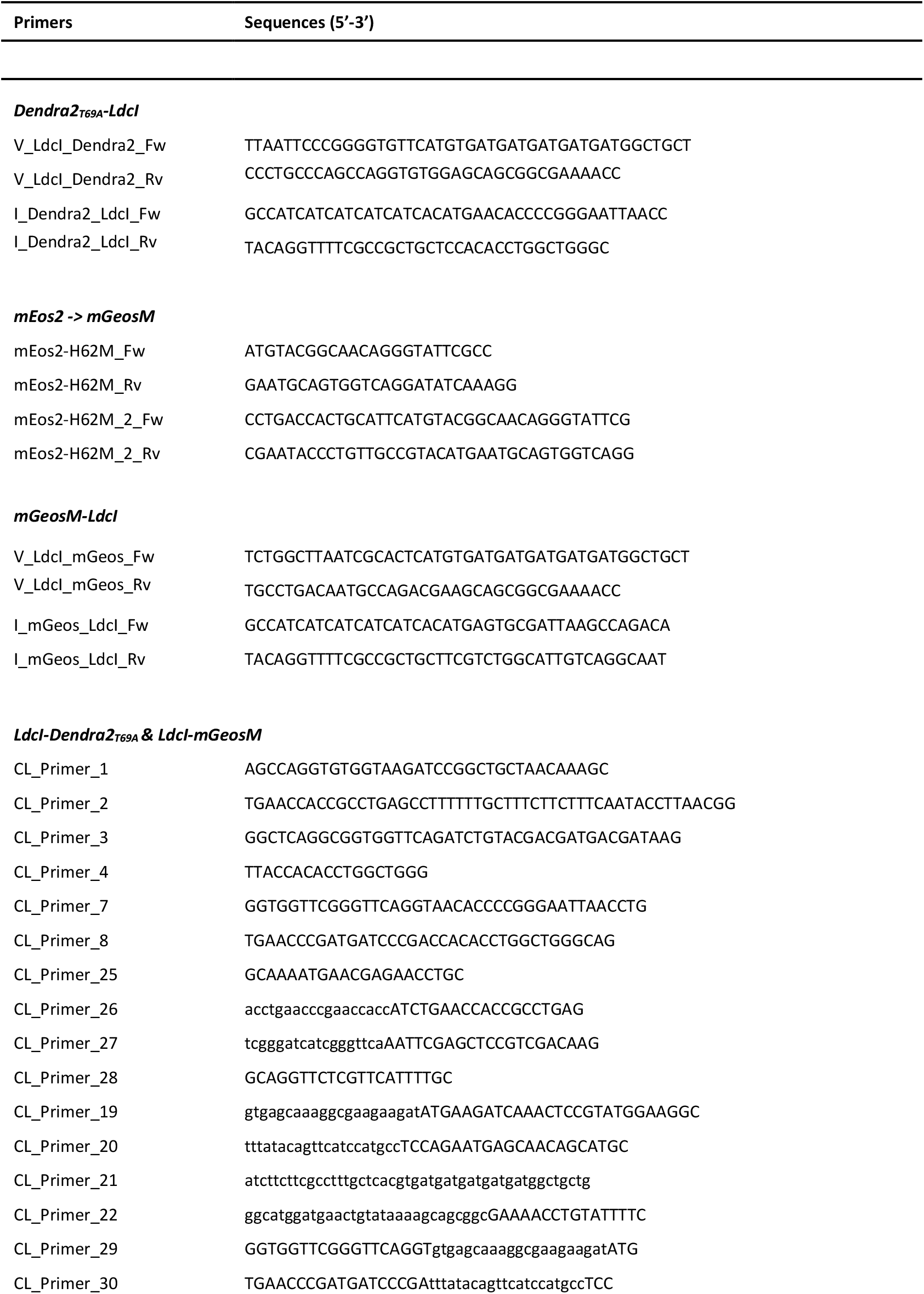

**Supplementary Table 4, C.**
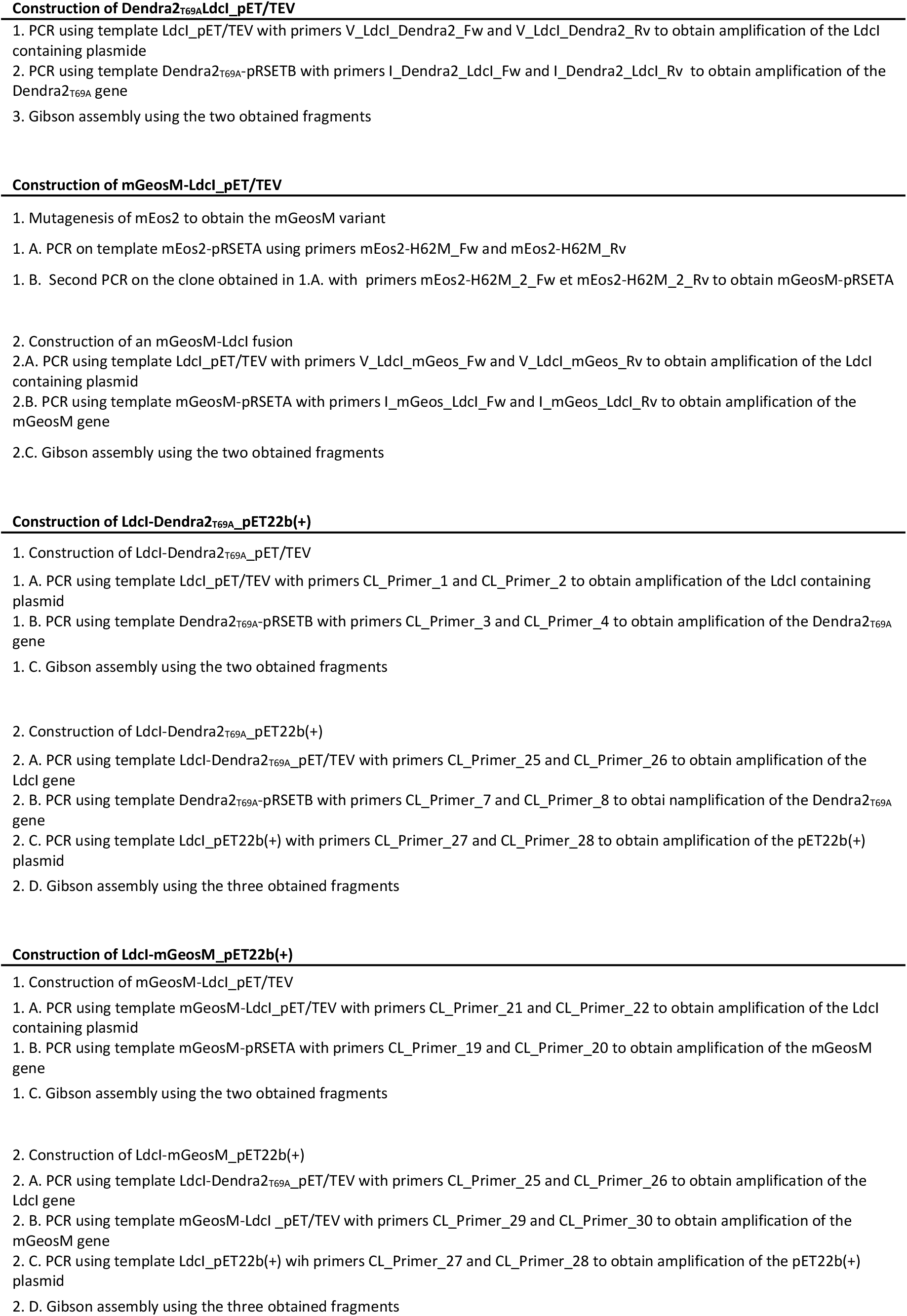

